# Sparse Coding in Temporal Association Cortex Improves Complex Sound Discriminability

**DOI:** 10.1101/2020.12.09.417303

**Authors:** L Feigin, G Tasaka, I Maor, A Mizrahi

## Abstract

The mouse auditory cortex is comprised of several auditory fields spanning the dorso-ventral axis of the temporal lobe. The ventral most auditory field is the temporal association cortex (TeA), which remains largely unstudied. Using Neuropixels probes, we simultaneously recorded from primary auditory cortex (AUDp), secondary auditory cortex (AUDv) and TeA, characterizing neuronal responses to pure tones and frequency modulated (FM) sweeps in awake head-restrained mice. As compared to primary and secondary auditory cortices, single unit responses to pure tones in TeA were sparser, delayed and prolonged. Responses to FMs were also sparser. Population analysis showed that the sparser responses in TeA render it less sensitive to pure tones, yet more sensitive to FMs. When characterizing responses to pure tones under anesthesia, the distinct signature of TeA was changed considerably as compared to that in awake mice, implying that responses in TeA are strongly modulated by non-feedforward connections. Together with the known connectivity profile of TeA, these findings suggest that sparse representation of sounds in TeA supports selectivity to higher-order features of sounds and more complex auditory computations.

## Introduction

The mouse auditory cortex is comprised of several auditory fields, the definition of which may vary between parcellation methodologies (Tsukano et al. 2016; Geissler, Schmidt, and Ehret 2016). The most ventral region of these fields is the auditory temporal association cortex, also known as auditory TeA. While both primary (AUDp) and secondary/ventral (AUDv) auditory fields were subject to significant research over several decades and in several species (Maor, Shalev, and Mizrahi 2016; Rothschild, Nelken, and Mizrahi 2010; Issa et al. 2014; Ohga et al. 2018; Issa et al. 2017), TeA remained largely unexplored.

In the rodent brain, TeA is a narrow and elongated cortical field, located below somatosensory, auditory and visual cortices. TeA is subdivided to these sensory modalities according to its corresponding positions along the rostro-caudal axis (Ramesh et al. 2018; Zingg et al. 2014; Yamashita et al. 2018). The auditory TeA region in the mouse or analogous fields in the rat were shown to be tonotopically organized and involved in auditory fear-conditioning (Dalmay et al. 2019; Quirk, Armony, and LeDoux 1997; Polley et al. 2007; Romero et al. 2020). Recently, the auditory TeA was shown to play a key role in maternal behavior in the context of the mother’s processing of pup ultrasonic vocalizations (Tasaka et al. 2020). Otherwise, to the best of our knowledge, little is known about the function or physiology of TeA.

Here, we set out to study basic response profiles of single neurons and populations in auditory TeA (hereafter, called TeA) of awake head-restrained mice. Using Neuropixels probes, (Jun et al. 2017), we recorded single unit activity from multiple neurons and cortical regions simultaneously. We used both pure tones and frequency-modulated (FM) sounds as auditory stimuli, while simultaneously recording responses in AUDp, AUDv and TeA. We describe the activity in TeA with respect to its upstream primary cortices. We found that, as compared to more primary stations, TeA exhibits sparser auditory-driven activity with more complex late activity that is likely modulated by non-feed-forward signals. Comparing to AUDp, TeA shows lower discriminability to simple tones, and better discriminability to the more complex FM sweeps. We also compared the response profile of these three cortices in two physiological states, wakefulness and anesthesia, which together show that TeA is a higher-order auditory processing station. TeA’s unique responses along the auditory hierarchy improves the computations of higher order sound features whilst coding of lower order sound features is compromised.

## Results

### Simultaneous recording from three auditory cortices in awake mice

To study the nature of auditory responses in TeA, we recorded extracellular spiking activity in response to sounds using the high-density probe, Neuropixels (Fig. 1a, Jun et al. 2017). We assessed responses in TeA, in head-fixed awake mice, with reference to auditory responses in two well-studied auditory cortices - primary auditory cortex (AUDp) and secondary/ventral auditory cortex (AUDv). To do so, we penetrated the brain with a single probe such that it diagonally traversed all three cortical regions; from dorsal to ventral: AUDp →AUDv→TeA (Fig. 1b). To validate the locations of our recordings we reconstructed (postmortem) the probe tracts, which were coated with a fluorescent lipophilic dye. We used DiI- and/or DiO-coated probes for multiple (up to three) sequential penetrations in each mouse (Fig. 1c; Methods). All probe trajectories were aligned to the Allen Brain Atlas coordinate framework validating the exact positions of our recordings in all mice (Fig. 1d, Shamash et al. 2018). In this manner, we obtained simultaneous recordings from all three auditory cortices in all mice (see Fig. S1 for precise locations and layer information of all probes). A representative 1.7-sec snippet of raw data recorded from AUDp →AUDv→TeA is shown in figure 1e. Following spike sorting, we obtained both well-isolated single units (SUs) and multi units (MUs). In total, we recorded from 12 probe penetrations in 5 mice, obtaining 1006 units (555 SUs and 451 MUs). Throughout the paper we present data from the well isolated SUs unless mentioned otherwise (AUDp, n=240 SU; AUDv, n=218 SU; TeA, n=97 SU; see Tables 1 and S1 for more details). Results from the MU data were pooled separately, and used only for specific analyses (Tables S1-S3).

**Figure 1.**
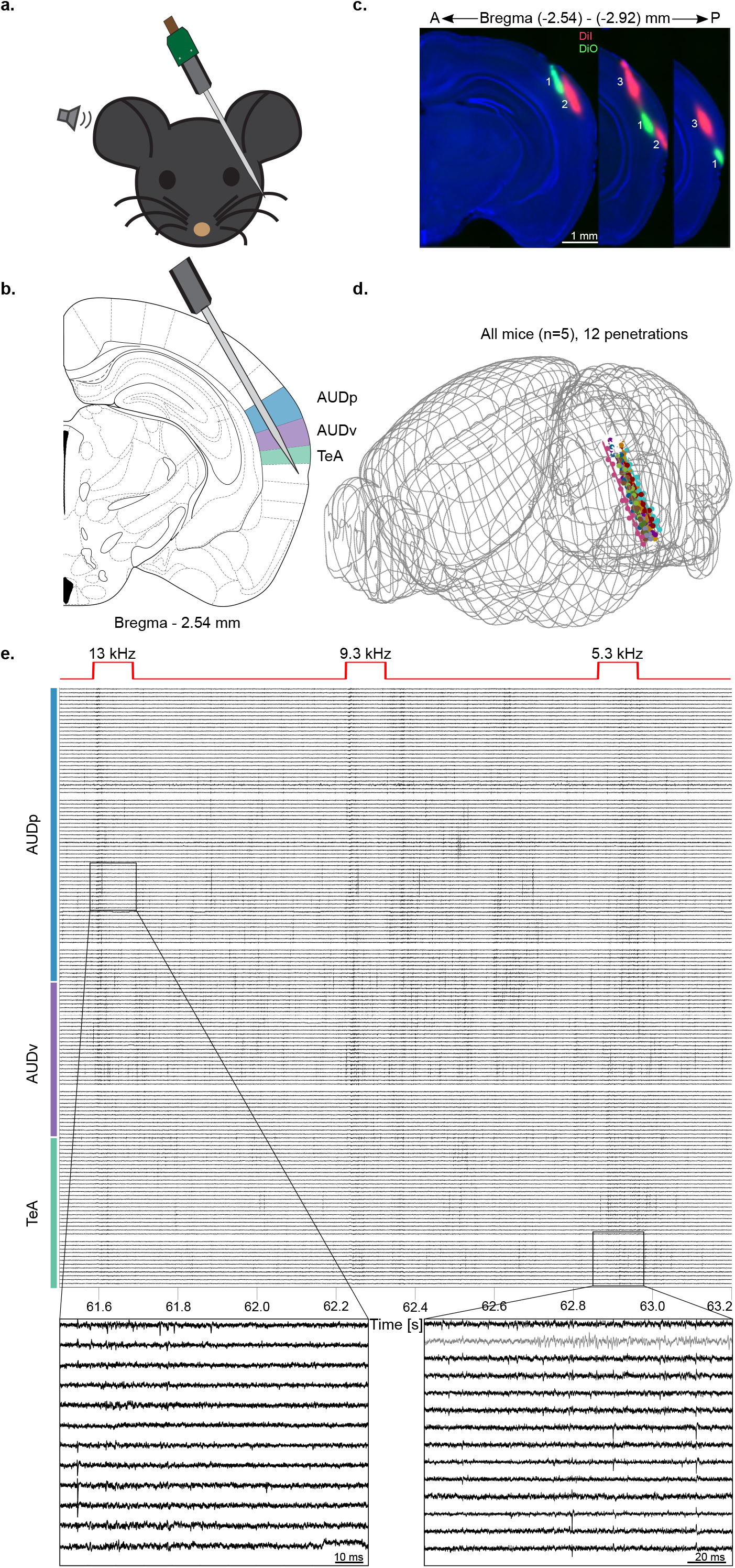
Simultaneous recording from three auditory cortices in awake mice. (a) Schematic representation of the experimental setup. (b) Probe penetration setting enabling simultaneous recording of activity from three auditory cortices (AUDp, AUDv and TeA; highlighted in color). (c) Three consecutive fluorescent images of coronal brain slices showing three probe tracks in one mouse. (d) Reconstructed trajectories of all probe tracts (12 penetrations, n=5 mice). (e) A 1.7sec snippet of the raw voltage traces recorded during pure tone presentation (total of 156 contacts shown). Top red trace shows the delivered sound stimuli, with captions indicating frequency in kHz. Color bar shows annotated regions according to channel and depth. Two activity snippets are shown in zoomed-in version (gray trace - internal reference channel).

### Auditory responses to pure tones in TeA become sparser

As expected from cortical responses in awake mice, spiking activity was highly heterogeneous in all of the three auditory regions (Fig. 2a; Montijn, Goltstein, and Pennartz 2015; Cembrowski and Spruston 2019; Zhan and Luo 2010; Tao et al. 2017). The first striking difference between TeA and the two other cortices was its sparser spiking activity. Spontaneous firing rates (FRs) decreased along the dorsal-to-ventral (D-V) cortical axis (Fig. 2b), all of which were distributed log-normally (Fig. S2). To characterize evoked auditory responses to pure tones, we presented thirty 100-msec pure tones with frequencies ranging between 3 to 80 kHz, each at 4 different attenuations (from 10-40 dB SPL). The vast majority of SUs in all three auditory cortices were significantly responsive to pure tones, with responses being either excitatory or suppressive (Table 1; see Methods for defining excitatory and suppressive responses). Excitatory responses to pure tones were more dominant than suppressive responses (Fig. 2c). However, while the fraction of excited units (out of all auditory units) remained high and consistent across all cortices, the fraction of units undergoing significant suppression decreased along the auditory D-V axis (Fig. 2c, from 68% to 49%).

**Figure 2.**
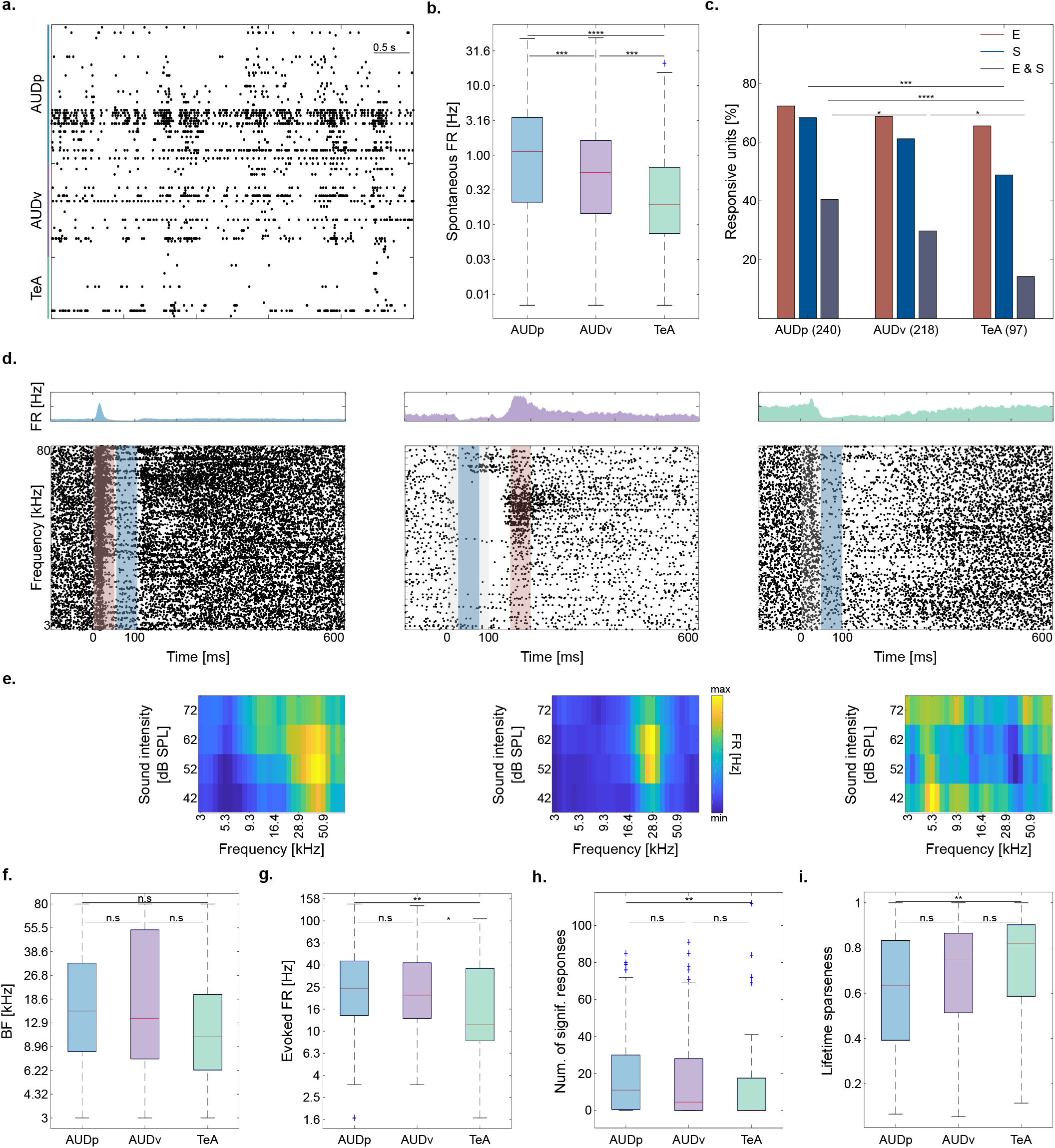
Pure tone response properties in TeA, AUDp and AUDv. (a) Raster plot showing 5 seconds of spontaneous activity. Color bar shows annotated regions according to channel and depth. (b) Spontaneous firing rates (log-scaled) of SUs in the three cortices (Median (IQR): AUDp-1.13 (0.21-3.48) n=240, AUDv-0.56 (0.14-1.63) n=218, TeA-0.19 (0.07-0.67) n=97). Spontaneous firing rates decrease along the auditory D-V axis (H(2)=35.68, p<1e-4; Kruskal-Wallis), are highest in AUDp, and lowest in TeA (AUDp vs. AUDv p=0.003, AUDp vs. TeA p<1e-4, AUDv vs. TeA p=0.003; TK HSD test). (c) Bar graph showing the distribution of response types to pure tones (E-excitatory, S-suppressive, E&S-both) in all cortical regions. Percentage computed out of all responsive units (number of SU shown in parentheses). Excitatory responses are abundant and their fraction is maintained high in all three cortices (H(2)=1.5,p=0.47; Kruskal-Wallis), while abundance of suppressed units (H(2)=10.06,p=0.006; Kruskal-Wallis. AUDp vs. AUDv-p=0.28 n.s; AUDp vs. TeA p<0.005; AUDv vs. TeA p=0.12 n.s; TK HSD test) as well as units showing both response types decreases from AUDp to TeA (H(2)=20.1, p<1e-4; Kruskal-Wallis; AUDp vs. AUDv-p=0.047; AUDp vs. TeA-p<1e-4; AUDv vs. TeA-p=0.03). (d) Three examples of SU responses to pure tones recorded in AUDp (left), AUDv (center) and TeA (right). Above each raster is the unit’s PSTH (scale adjusted per unit). Stimulus is presented between 0-100 msec (gray shaded region), and each units’ maximal or minimal response window is marked by a red and/or blue bar, respectively. The units are categorized as Excitatory & Suppressed (left and center), and suppressed (right). (e) FRAs corresponding to the three SUs shown in ‘d’. FRAs pixel values correspond to the excitatory (left and center) or suppressive (right) response window. (f) SU best frequencies across brain regions (Median (IQR): AUDp-15.5 (8.3-32.3) kHz, n=164 SUs; AUDv-13.8 (7.4-53.9) kHz, n=136 SUs; TeA-10.4 (6.3-20.0) kHz, n=55 SUs). We found no effect of brain region over BF (H(2)=3.07, p=0.21; Kruskal-Wallis), as well as no difference between BF medians (AUDp vs. AUDv-U=2.470e4, p=0.98 n.s; AUDp vs. TeA U=1.875e4, p=0.08 n.s; AUDv vs. TeA U=1.357e4, p=0.14 n.s; Wilcoxon ranksum test). (g) Evoked firing rates at BFs (Median (IQR): AUDp-24.5 (13.9-43.3) Hz, n=164 SUs; AUDv-21.2 (13.1-41.7) Hz, n=136 SUs; TeA-11.4 (8.2-37.2) Hz, n=55 SUs). SUs in TeA had lower FRs compared to units from both AUDp and AUDv (H(2)=11.13, p=0.004; Kruskal-Wallis. AUDp vs. AUDv-p=0.63 n.s; AUDp vs. TeA-p=0.002; AUDv vs. TeA-p=0.03; TK HSD test). (h) Bandwidth of SUs assessed by the number of (frequency x attenuation) combinations evoking a significant response (Median (IQR): AUDp-11 (0.530), n=164 SUs; AUDv-4.5 (0-28), n=136 SUs; TeA-0 (0-17.5), n=55 SUs). Bandwidth in TeA was smaller than in AUDp (H(2)=11.70, p=0.003; Kruskal-Wallis. AUDp vs. AUDv-p=0.17 n.s, AUDp vs. TeA-p=0.004; AUDv vs. TeA-p=0.25 n.s; TK HSD test). (i) Lifetime sparseness of SUs (Median (IQR): AUDp-0.64 (0.39-0.83), AUDv-0.75 (0.51-0.87), TeA-0.82 (0.59-0.90)). Sparseness was larger in TeA compared to AUDp (H(2)=11.32, p=0.003; Kruskal-Wallis. AUDp vs. AUDv – p= 0.08 n.s; AUDp vs. TeA p=0.004; AUDv vs. TeA p=0.3 n.s; TK HSD test).

The rich nature of pure tone responses is poorly captured when being described as excitatory or suppressive alone. For example, some SUs responded by interleaved excitation and suppression (Fig. 2d, left and center), while others responded to different frequency bands with “on” or “off” or both response types (Fig. S3a). Excitatory responses were highly diverse; from being transient and time-locked to the stimulus (Fig. S3b), through jittery spiking responses (Fig. S3c), and up to strong and long-lasting activity (Fig. S3d; see Fig. S3e-h for more examples). Suppressive responses were as rich and heterogenous as the excitatory responses (see Fig. 2d; Fig. S3i). Frequency response areas (FRAs) were heterogeneous in all regions, ranging from V-shaped FRAs (Fig. 2e-left), I-shaped (Fig. 2e-center) or having more complex tuning properties (e.g., Fig 2e-right). This observation is consistent with previous reports on AUDp response heterogeneity, extending it to both TeA and AUDv (Rothschild, Nelken, and Mizrahi 2010; Bowen, Winkowski, and Kanold 2020; Bandyopadhyay, Shamma, and Kanold 2010). In all cortices, the span of best frequencies (BFs) in our recordings was similarly widespread (Fig. 2f; Fig. S4a). Nonetheless, cross-region transitions (e.g. AUDp to AUDv or AUDv to TeA) were characterized by larger BF differences between neighboring SUs compared to neighboring units within a single region, as expected from crossing anatomically-defined boundaries (Fig. S4b). Evoked firing rates were significantly higher in AUDp and AUDv as compared to TeA (Fig. 2g); all of which were also distributed log-normally (Fig. S4c).

Next, we examined the breadth of excitatory response profiles in more detail employing three parameters of sparseness - broadness of tuning, population sparseness and lifetime sparseness. For assessing the broadness of tuning of each unit we examined the number of frequency x attenuation combinations evoking a statistically significant response (Fig. 2h). The average tuning of units in TeA was narrower as compared to units in AUDp (Fig. 2h). In fact, the majority of SUs in TeA were only weakly responsive and not strongly tuned (e.g., 54% of SUs had no significant responses in their FRA). We estimated population sparseness (also referred to as activity sparseness by Willmore and Tolhurst 2001), by calculating the fraction of SUs responsive to each stimulus. In TeA, the response of the population was sparser as compared to AUDp and AUDv (Fig. S5a,b). Finally, we calculated lifetime sparseness, a statistic describing how dispersed is a neurons’ firing across different stimuli. Lifetime sparseness was significantly larger in TeA as compared to AUDp (Fig. 2i). To conclude, our data show that representation of frequency in SUs of awake mice becomes sparser along the dorso-ventral axis of the auditory cortex.

### TeA responses to pure tones are delayed and protracted

Recently, using monosynaptic rabies tracing, we showed that TeA is one synapse downstream of AUDp (Tasaka et al. 2020). We, therefore, expected that auditory responses in TeA will lag behind AUDp by several milliseconds. A units’ latency can be estimated in several different ways, each of which capturing different aspects of the neuronal response. Thus, we used two separate measures for describing unit latency - ‘minimal latency’ and ‘latency to peak’ (Fig. 3a). To calculate minimal latency, we isolated solely excited units with response windows that were detected up to 110 ms from stimulus onset (and effectively 10 ms post stimulus offset, to avoid offset responses). In these units, we calculated the median latency to the 1^st^ spike in the response window at the unit’s BF. As we anticipated, the minimal latency in TeA was larger as compared to AUDp (Fig. 3b; Δ=14 ms). To calculate latency to peak, we measured the latency to the extremum of each units’ response. This statistic was measured for units that were excited (as latency to peak) as well as for units that were suppressed (as latency to minima). Latency to peak, but not to minima, was 49 ms higher in TeA as compared to AUDp (Fig. 3c and d, respectively). The time to peak of individual units was diverse, tiling a wide temporal window (from 7-523 ms; Fig. 3e). As compared to AUDp, the distribution of peak times in TeA was shifted to slower values (Fig. 3e). Notably, and in agreement with previous reports (Mormann et al. 2008), latency and sparseness were positively correlated (Fig. S5c).

**Figure 3.**
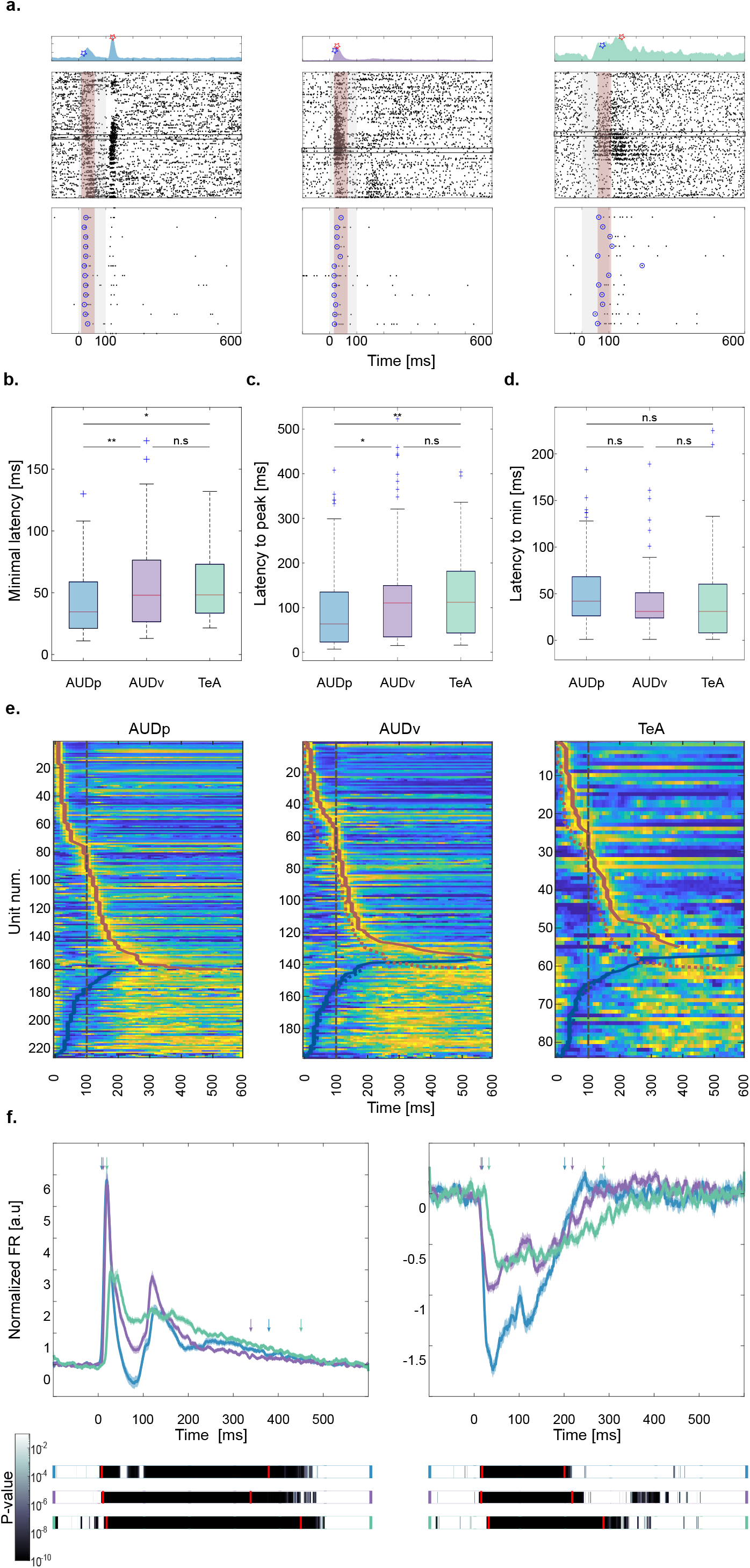
Temporal response profile of SUs in TeA, AUDp, and AUDv. (a) Three representative example SUs from AUDp (left), AUDv (center) and TeA (right). Top: PSTHs showing the units’ minimal latency (blue marker), and latency to peak (red marker). Center: raster plots (details as in Fig. 2d). Red shaded regions mark the 50 ms maximal early response windows of each unit, and black rectangles mark the units’ early BF trials. Bottom: Magnification of early BF trials with circles surrounding the first spike in each trial. (b) Minimal latency across brain regions (Median (IQR): AUDp–34 (21-59) ms n=123, AUDv-48 (26-76) ms n=101, TeA-48 (33-73) ms n=38). Minimal latency in AUDp was shorter compared to both other auditory cortices (H(2)=12.43, p=0.002; Kruskal-Wallis. AUDp vs. AUDv-p=0.007; AUDp vs. TeA-p=0.02; AUDv vs. TeA-p=0.89 n.s; TK HSD test). (c) Latency to peak across brain regions (Median (IQR): AUDp-63 (23-135) ms n=164, AUDv-110 (34-149) ms n=136, TeA-112 (43-181) ms n=55). TeA units’ latency to peak was larger than that of AUDp units (H(2)=12.48, p=0.002; Kruskal-Wallis. AUDp vs. AUDv-p=0.01; AUDp vs. TeA-p=0.007; AUDv vs. TeA-p=0.6 n.s; TK HSD test). (d) Latency to minima across brain regions (Median (IQR): AUDp-42 (26-68) ms n=63, AUDv-31 (24-51) ms n=62, TeA-31 (8-60) ms n=29). Time to minima was similar across cortices (H(2)=1.78, p=0.41 n.s; Kruskal-Wallis). (e) Heat maps of SU FRs in AUDp (left), AUDv (center) and TeA (right) sorted by latency to peak for excited units (top to bottom) and latency to minima for suppressed units (bottom up). Thick red and blue lines mark times to peak and to minima, respectively. The trajectory of AUDp is overlaid on top of the heat maps of AUDv and TeA (dashed colored lines). Time of stimulus offset is marked with gray dashed lines. (f) Normalized population response averaged across pure tone stimuli for excited (left) and suppressed (right) SUs. Bold lines and shaded regions mark mean response and SEM, respectively (blue-AUDp, purple-AUDv, green-TeA). Bars (bottom) show the p-value of the instantaneous activity vs baseline for each region. The time of deviation from baseline was larger for TeA (Excited: AUDp-5 ms, AUDv–8 ms, TeA-16 ms; Suppressed: AUDp-15 ms, AUDv-13 ms, TeA-30 ms; t-test Bonferroni corrected for MC over time bins to s.t p(thresh)=7.1e-5). Duration of responses in TeA was more than 69 ms longer than in other regions (Time to return to baseline, Excited: AUDp-375 ms, AUDv-335 ms, TeA-446 ms; Suppressed: AUDp-201 ms, AUDv-215 ms, TeA-284 ms, paired t-test corrected for MC s.t p(thresh)=7.1e-5). Time to deviate and return to baseline is marked by red lines and colored arrows.

The normalized population response of all excited units (n: AUDp=164 SUs, AUDv=132 SUs, TeA=55 SUs) across all stimuli further shows that TeA activity arises later than both AUDp and AUDv (Fig. 3f). Moreover, population activity in TeA was not only delayed, but was also more persistent and decayed with slower time scales as compared to AUDp and AUDv. It takes the population response in TeA at least 70 ms longer than in AUDp and AUDv to return to baseline (Fig. 3f). We conclude, therefore, that TeA exhibits delayed and prolonged auditory evoked responses as compared to both AUDp and AUDv.

### TeA responses to FMs are excitatory and sparse

While pure tones are the fundamental elements of any sound, they lack the rich spectro-temporal attributes of more natural sounds. To obtain a more complete characterization of basic auditory responses in TeA, we used frequency modulated sounds (FMs). Specifically, we used linear and logarithmic FM sweeps (Fig. 4a; note that only the logarithmic FM stimulus set is shown). The FM stimuli set consisted of 10 sweeps of each type (linear/logarithmic), all spanning the range between 4-64 kHz at varying slopes, and presented at a sound intensity of 62 dB SPL. For every FM type, 5 stimuli were rising (from 4 to 64, gradually) and 5 were falling in the exact opposite direction. Different stimuli had different durations, from 40 ms up to 2 seconds (Fig. 4a).

**Figure 4.**
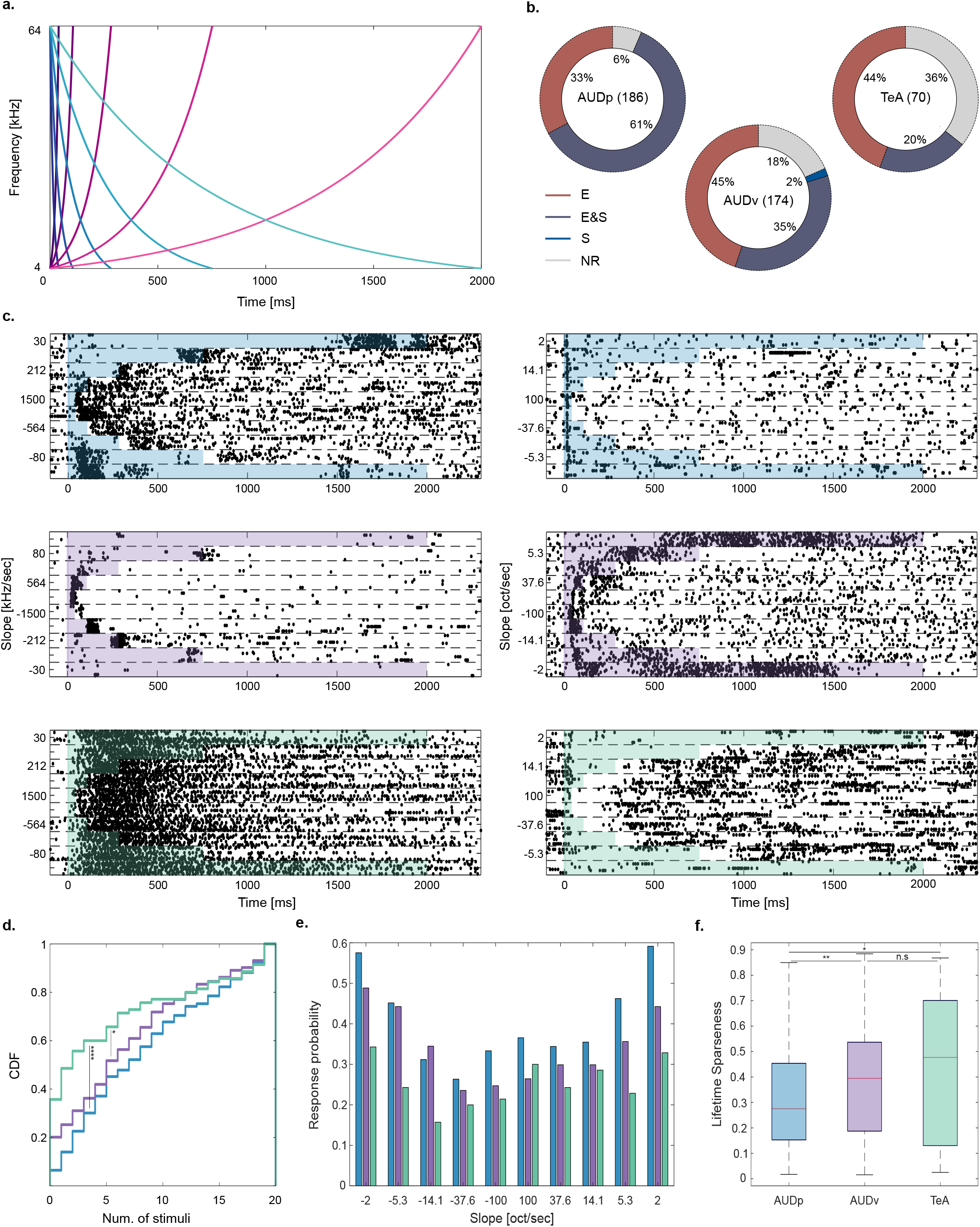
Basic response to FMs in TeA, AUDp, and AUDv. (a) FM stimulus set. Frequency *vs.* time representations of rising (pink palette) and falling (blue palette) logarithmic FM sweeps. Darker colors correspond to larger modulation speeds and shorter stimuli. (b) Donut plots summarizing FM response types. E (excited), E & S (excited and suppressed), S (suppressed) and NR (not responding). The number of SUs recorded in region are shown in parentheses. In all regions most SUs exhibit excitatory responses, with the fraction of excited SUs decreasing along the D-V axis of the cortex (Fraction excited SUs: AUDp–174/186=0.93, AUDv-139/174=0.80, TeA-45/70=0.64; H(2)=33.54 p<1e-4; Kruskal-Wallis. AUDp vs.TeA p<1e-4, AUDp vs.AUDv p=0.001, AUDv vs.TeA p=0.009; TK HSD test). (c) Representative examples of SU FM responses from AUDp (top row), AUDv (central row) and TeA (bottom row). Left column shows responses to linear FM sweeps, while the right column shows responses to logarithmic FM sweeps. Shaded regions mark the stimulus durations. (d) CDF of the number of FM stimuli evoking significant excitatory response in SUs of AUDp (blue) AUDv (purple) and TeA (green). Responsivity was lower in TeA compared to both AUDp and AUDv (H(2)=18.20 p=0.0001, Kruskal-Wallis. AUDp vs. AUDv-p=0.07 n.s, AUDp vs. TeA-p<1e-4, AUDv vs. TeA-p=0.03; TK HSD test). (e) Population sparseness presented as the response probability of SUs in each region to logarithmic FMs. (f) Lifetime sparseness calculated for all FM stimuli (Median (IQR): AUDp-0.27(0.15-0.45) n=174, AUDv-0.39 (0.19-0.54) n=139, TeA-0.48 (0.13-0.70) n=45). Sparseness was lower in AUDp compared to both other cortices (H(2)=12.84 p=0.002; Kruskal-Wallis. AUDp vs. AUDv-p=0.009, AUDp vs. TeA-p=0.01, AUDv vs. TeA p=0.7 n.s; TK HSD test).

The majority of units in all regions exhibited significant responses to at least one FM stimulus (Fig. 4b; fraction non-responsive SUs: AUDp-12/186=0.06, AUDv-32/174=0.18, TeA-25/70=0.36). We focused our analysis solely on FM responses that were excitatory since they were the vast majority (Fig. 4b). On average, responses to the different FM types (linear and logarithmic) were qualitatively and quantitatively similar (Fig. S6). Thus, unless stated otherwise, we pooled the data from the two stimulus sets. SU responses to FMs were diverse in their nature in all three cortical regions (Fig. 4c). Some units had excitatory responses during specific time windows along the sweep, corresponding to the units preferred frequency range (Fig. 4c; center left and right). Other response types could be suppressed during all or most sound duration, with potential excitatory overshoots at specific stimulus times (Fig. 4c; left: top, right: bottom). Some units responded to sound onset, independent of the FM type and direction (Fig. 4c; right: top), and a small fraction of the SUs was characterized by persistent firing (Fig. 4c; left: bottom). More examples showing the high diversity of responses to FM sounds are shown in Fig. S7.

Next, we examined the sparseness of FM responses with similar measures to those we used for pure tone responses. We assessed the broadness of each units’ FM tuning by counting the number of FM stimuli evoking a significant response. Compared to both AUDp and AUDv, TeA SUs were sparser, and on a per-unit basis responded to fewer FM stimuli (Fig. 4d). To assess population sparseness, we counted the fraction of SUs in each region having a significant response to each FM stimulus. Here again, the number of TeA units being activated by FM stimuli was consistently smaller compared to AUDp and AUDv (Fig. 4e; fraction of population responsive: TeA<AUDp - 20/20 FM stimuli, TeA<AUDv 15/20 FM stimuli, AUDv<AUDp - 18/20 FM stimuli). Finally, we calculated the lifetime sparseness of FM responses. Sparseness was larger for TeA and AUDv units compared to AUDp, with TeA units exhibiting a relatively wide distribution of sparseness values (Fig. 4f). Therefore, and qualitatively similar to the pure tone analysis, TeA responses to FMs becomes sparser along the D-V cortical axis.

### FM-direction and FM-speed selectivity are reduced in TeA

One way to characterize FM responses is to assess direction and speed selectivity (Nelken and Versnel 2000; Zhang et al. 2003; Sollini et al. 2018; Trujillo, Carrasco, and Razak 2013). We calculated direction selectivity index per SU as 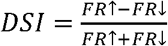 (see Methods). A DSI of 1 indicates that the unit’s response it strictly to the rising FMs and *vice versa* for a DSI of −1. A DSI of 0 indicates no direction selectivity. Unlike AUDp and AUDv, SUs in TeA had an average DSI different and larger than zero, indicating a slight preference for rising FMs (Fig. 5a). In some SUs, it was evident that response sparseness was accompanied by strong FM direction selectivity (Fig. 5b). Indeed, DSI and lifetime sparseness were strongly and significantly correlated (Fig. 5c; r=0.602 p<1e-4), suggesting that TeA’s sparseness contributed to its FM selectivity.

**Figure 5.**
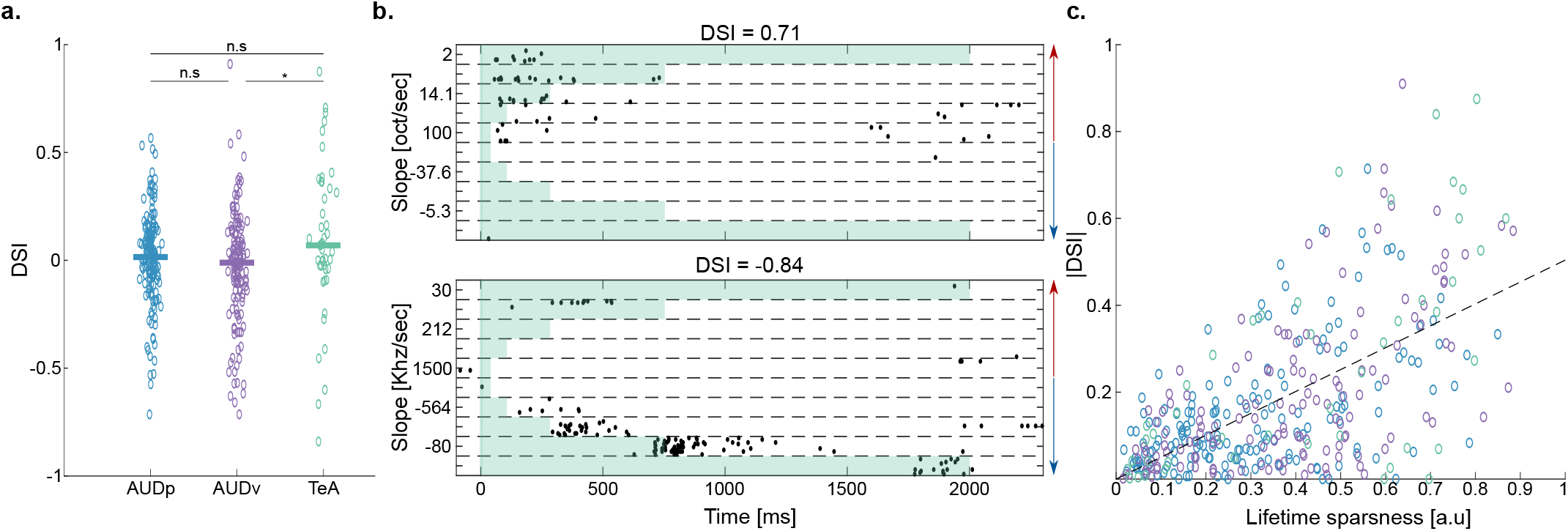
Selectivity to FMs in TeA, AUDp, and AUDv. (a) Scatter plot of direction selectivity index (DSI) values. Median values marked by bold lines (Median (IQR): AUDp-0.015 (−0.075-0.123) n=174, AUDv- −0.011 (−0.143-0.127) n=142, TeA-0.069 (−0.040-0.341 n=45). DSI was different from zero only in TeA (AUDp-z=1.29, p=0.20; AUDv-z=-1.05, p=0.29; TeA-z=1.98, p=0.048; Wilcoxon signed-rank test), with TeA SUs showing a greater preference towards rising FMs (H(2)=7.11 p=0.029, Kruskal-Wallis. AUDp vs. AUDv-p=0.31 n.s, AUDp vs. TeA-p=0.21 n.s, AUDv vs. TeA-p=0.024; TK HSD test). (b) Raster plots of two SUs from TeA exhibiting high selectivity to rising (top) and falling (bottom) FM stimuli, respectively. Red and blue arrows mark FM direction (rising and falling, respectively). Each unit’s DSI is noted above the raster. Rasters correspond to logarithmic (top) and linear (bottom) FM stimuli. (c) Scatter plot of absolute DSI values vs. lifetime sparseness. Units are color-coded according to their cortical origin. Black dashed line shows fitted regression line between the two parameters (|DSI|=0.48*LF+0.01, R^2^=0.362).

To examine speed-selectivity we plotted response magnitudes to the different FM slopes (Fig. Fig. S8a). In AUDp and AUDv response magnitudes were highest for low speed FMs and decreased as speeds grew (Fig. S8a-blue and magenta). In TeA this effect was attenuated, and it seemed short latency FMs were equally preferred (Fig. S8a - green). Since responses were calculated based on optimal response windows per FM which were all the same size (50 msec), we presume different FM durations were not the source of these variations, and they seem to reflect a genuine “bimodal” speed selectivity in TeA.

### Population responses in TeA differ in discriminability for pure tones and FM sweeps

While single neurons are the fundamental building blocks of sensory coding, sensory codes are likely read as a population. Moreover, correlations among neurons have been suggested as key features of the population neural code (Averbeck, Latham, and Pouget 2006; Nirenberg and Latham 2003; M. R. Cohen and Kohn 2011). Thus, we next characterized pairwise correlations of simultaneously recorded SUs. For simplicity we focus on two brain regions only – AUDp and TeA, and use responses to pure tones. First, we assessed pairwise signal correlations (SC), which describes the similarity in FRA tuning (Gawne and Richmond 1993; M. R. Cohen and Kohn 2011). The mean SC within regions were generally low in both regions (Fig. 6a), yet significantly larger than expected from shuffled data (p<1e-4 for all comparisons of shuffled vs data; Kolmogorov-Smirnoff test). Cross region correlations were even lower which is expected because correlations are known to decay with increasing distance (Fig. 6a; Rothschild, Nelken, and Mizrahi 2010; Smith and Kohn 2008). Nonetheless, highly correlated pairs were evident in all distributions (Fig. 6a; see positive “tails” in all distributions). Second, we measured noise correlations (NC), the trial-to-trial tendency of units firing rates to fluctuate together (Averbeck, Latham, and Pouget 2006; M. R. Cohen and Kohn 2011). Mean NC were centered around 0, although within-region correlations significantly higher than cross region correlations (Fig. 6b). Positive tails of higher correlated units were evident for NC as well (Fig. 6b). Finally, we tested the relationship between SC and NC (Rothschild, Nelken, and Mizrahi 2010; Smith and Kohn 2008). SC and NC were positively correlated, with higher interaction within regions as compared to among cortical regions (Fig. 6c; AUDp-r=0.396 p<1e-4, TeA-r=0.444 p<1e-4, AUDp-TeA-r=0.227 p<1e-4). These data show that SC and NC behave largely similarly along the auditory cortical hierarchy.

**Figure 6.**
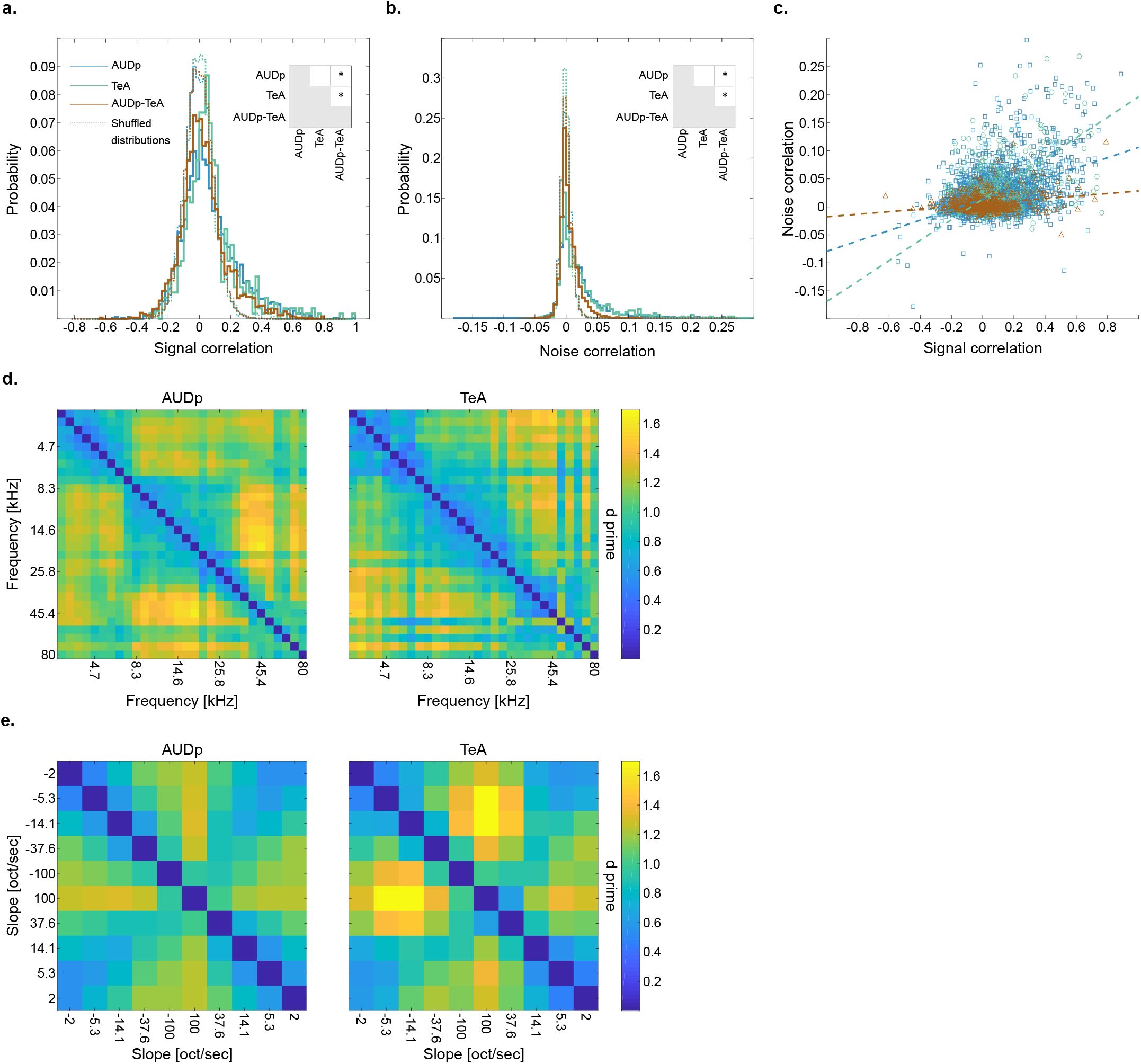
Population analysis of pure tones and FMs in AUDp and TeA. Pairwise signal correlations (SC) in AUDp (blue), TeA (green) and between both regions (AUDp-TeA, brown) (Median (IQR): AUDp-0.04 (-0.05-0.16) n=3042, TeA-0.05 (−0.02-0.13) n=529, AUDp-TeA-0.01 (−0.06-0.09) n=2296). Correlation in AUDp and TeA were similar and larger than the cross region correlations (AUDp vs. TeA - U(3042,529)=5.406e6 p=0.22, AUDp vs. cross-region - U(3042,2296)=8.549e6 p<1e-4, TeA vs. cross-region - U(529,2296)=8.494e5 p<1e-4; Wilcoxon ranksum test. Significant comparisons shown in inset). Dashed lines represent shuffled distributions for each color-corresponding curve. All distributions were different from shuffled (p<1e-4 for all comparisons, Kolmogorov-Smirnoff test). (b) Same as (a) for noise correlations (NC; Median (IQR): AUDp-0.009 (-0.001-0.028) n=3042, TeA-0.011 (-0.001-0.034) n=529, AUDp-TeA - 0.001 (-0.003-0.011) n=2296). Here, too, correlations were indistinguishable between AUDp and TeA and larger in both than cross region correlations (AUDp vs. TeA - U(3042,529)=5.406e6 p=0.22, AUDp vs. crossreg - U(3042,2296)= 9.016e6 p<1e-4, TeA vs. cross-region - U(529,2296)=9.128e5 p<1e-4; Wilcoxon ranksum test). All distributions were different from shuffled (p<1e-4 for all comparisons, Kolmogorov-Smirnoff test). (c) Scatter plot showing relationship between noise and signal correlations. Dashed lines show linear fits for each individual population. Linear correlation between the two was strongest for TeA and weakest for cross region correlations (Linear fit, R^2^: AUDp-NC=0.09*SC+0.03 R^2^=0.157; TeA-NC=0.18*SC+0.01 R^2^=0.198; TeA-AUDp-NC=0.02*SC+0.005 R^2^=0.052). (d) Matrices of d primes for all pure tone pairs (62 dB SPL) calculated based on the activity of all SUs in AUDp (left, n=240 SUs) and TeA (right, n=97 SUs). Pairwise pure tone discriminability was larger in AUDp compared to TeA (mean±std: AUDp-1.06±0.23, TeA-0.98±0.26; t(435)=7.75 p<1e-4, paired sample t-test). (e) Matrices of d primes for logarithmic FM sweep pairs calculated based on the activity of all SUs in AUDp(left, n=186 SUs) and TeA (right, n=70 SUs). Pairwise FM sweep discriminability was larger in TeA compared to AUDp (mean±std: AUDp-0.93±0.23, TeA-1.01±0.32; t(45)=2.83 p=0.007, paired sample t-test).

Next, we set to see whether the neuronal populations in AUDp and TeA exhibit differences in their discriminability for both types of sounds we presented. We calculated pairwise d primes for population responses (see methods) to all pure tones presented at 62 dB SPL (Fig. 6d) and logarithmic FM sweeps (Fig. 6e). The population of neurons in AUDp showed significantly higher discriminability between pure tone pairs, as compared to TeA. The was reversed for discriminability between the different FM sweeps. In TeA, d-prime values of FM pairs were significantly higher than in AUDp. These data suggest that neurons in TeA are more prone to dissociate more complex auditory stimuli as compared to more primary cortices.

### Distinct effects of anesthesia on TeA and AUDp

Accumulating evidence suggest that anesthesia has a stronger effect on higher order cortices as compared to primary cortices (Mashour 2014; Jordan et al. 2013). Thus, since TeA is downstream of AUDp, we hypothesized it will be more strongly affected by anesthesia. To this end, we conducted a separate experiment, where we recorded in Ketamine-Medetomidine anesthetized mice, and assessed SU responses to pure tones only (6 probe penetrations in 5 anesthetized mice, obtaining 199 SUs from both regions; AUDp-n=151, TeA-n=48; See table S4 for details and layer information). Responsivity was significantly larger in the anesthetized compared to the awake state in both regions (AUDp-150/151=99% responsive SUs, TeA-47/48=98% responsive SUs; F(conscious state)=10.78 p=0.0011, F(brain region)=7.18 p=0.008; 2-way ANOVA). Indeed, anesthesia had a distinct signature on AUDp as compared to TeA. Spontaneous activity in TeA significantly increased, leveling up with spontaneous activity in AUDp, which did not change (Fig. 7a). Under anesthesia, evoked FRs were significantly lower in AUDp, and higher in TeA, such that responses were now stronger in TeA (Fig. 7b). Furthermore, the sparseness which was a prominent characteristic of TeA was strongly affected, resulting in similar levels of sparseness in both regions during anesthesia (Fig. 7 c,d).

**Figure 7.**
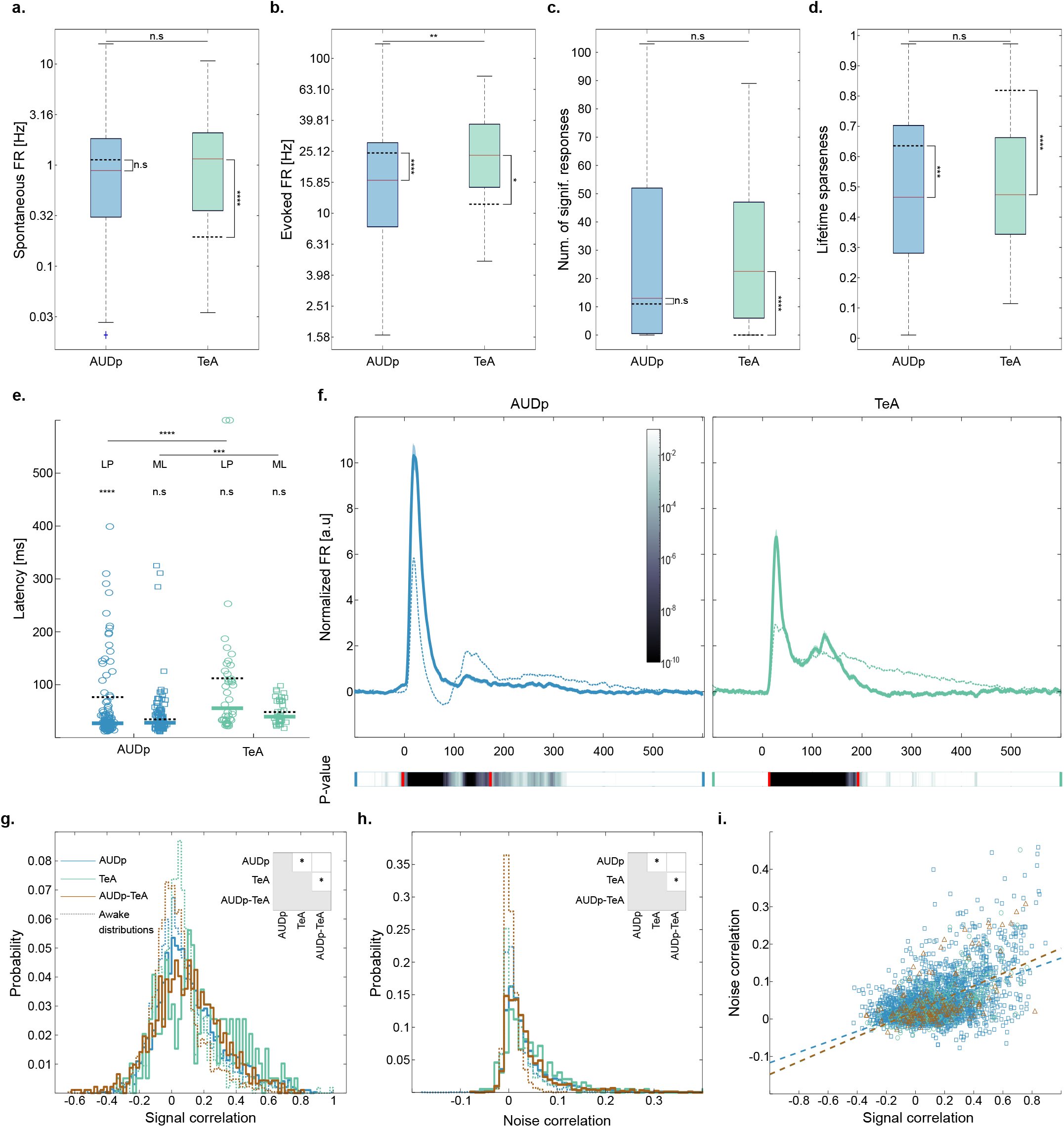
Anesthesia affects TeA and AUDp in distinct ways. (a) Spontaneous FRs in anesthetized AUDp and TeA (Median (IQR): AUDp – 0.88 (0.30-1.83) Hz n=151, TeA-1.15 (0.35-2.09) Hz n=48) are similar (U(151,48)=14704 p=0.25; Wilcoxon ranksum test). Anesthesia induces a significant increase in TeA spontaneous FR (AUDp-U(240,151)=49169 p=0.05 n.s; TeA-U(97,48)=6011 p<1e-4; Wilcoxon ranksum test Bonferroni corrected for MC). Dashed black lines mark awake population medians, adjacent asterisks mark within region anesthetized vs. awake comparisons. (b) Evoked firing rates in response to BFs (Median (IQR): 16.34 (8.17-28.59) Hz n= 132, TeA-23.69 (14.70-37.58) Hz n=42). Evoked FRs in the anesthetized cortex are higher in TeA compared to AUDp (U(132,42)=10752 p=0.005; Wilcoxon ranksum test). Anesthesia reduced evoked activity in AUDp and raised it in TeA (AUDp-U(164,132)=27472 p<1e-4, TeA-U(55,42)=2378 p=0.02; Wilcoxon ranksum test Bonferroni corrected for MC). (c) Bandwidth of SUs in the anesthetized cortex assessed by the number of (frequency x attenuation) combinations evoking a significant response (Median (IQR): AUDp-13 (0.5-52) n=132, TeA-22.5 (6-47) n=42). TeA bandwidth is larger under anesthesia compared with the awake state (AUDp-U(164,132)=27472 p=0.07 n.s; TeA-U(55,44)=2146 p<1e-4; Wilcoxon ranksum test Bonferroni corrected for MC), and is comparable with bandwidth in AUDp (U(132,42)=11321 p=0.42 n.s; Wilcoxon ranksum test). (d) Lifetime sparseness of SUs in the anesthetized cortex (Median (IQR): AUDp– 0.46 (0.28-0.70) n=132, TeA– 0.47 (0.34-0.66) n=42). Sparseness was similar across regions (U(132,42)=11423 p=0.65 n.s; Wilcoxon ranksum test) and lower compared to the awake state (AUDp– p= 0.0003, TeA-p<1e-4; Wilcoxon ranksum test Bonferroni corrected for MC). (e) SU minimal latencies (ML, right column, squares; Median (IQR): AUDp-28 (20-45) ms n=121, TeA-38 (30-70) ms n=37) and latencies to peak (LP, left column, circles: Median (IQR): AUDp-27 (21-24) ms n=132, TeA-55 (33-120) ms n=42) in the anesthetized state. Median values are marked with thick lines. Awake median values are marked with dashed black lines. Both latency parameters were larger in TeA (Minimal latency: U(121,37)=8785 p=0.0006; Latency to peak: U(132,42)=10258 p<1e-4; Wilcoxon ranksum test), however on a per region basis tended to be shorter than for the awake condition (Minimal latency: AUDp-U(123,121)=27648 p<1e-4, TeA-U(38,37)=2988 p=0.03 n.s; Latency to peak: AUDp-U(164,132)=15955 p=0.11 n.s, TeA-U(55,42)=1518 p=0.43 n.s; Wilcoxon ranksum test Bonferroni corrected for MC). (f) Normalized population response averaged across pure tone stimuli for excited SUs (AUDp-n=132, TeA=42) under anesthesia. Bold lines and shaded regions mark mean response and SEM respectively (left, blue-AUDp; right, green-TeA). Population responses from the awake condition are shown as dashed lines for comparison. Bottom color bars show p-value for comparisons of instantaneous activity to baseline, red lines marking beginning (left) and end (right) of deviation from baseline. Return to baseline activity (AUDp-171 ms, TeA-192 ms; paired t-test corrected for MC over time bins) was shorter than during wakefulness for both cortices. (g) Pairwise signal correlations (SC) in AUDp (blue), TeA (green) and between both regions (AUDp-TeA, brown) under Ketamine-Domitor anesthesia (Median (IQR): AUDp-0.077 (−0.025-0.220) n=2915, TeA-00133 (-0.018-0.327) n=198, AUDp-TeA-0.097 (-0.034-0.238) n=918). Correlations were highest in TeA (AUDp vs. TeA – U(2915,196)=4.495e6 p=<1e-3, AUDp vs. cross-region – U(2915,918)=5.572e6 p=0. 7 n.s, TeA vs. cross-region – U(196,918)=1.215e5 p=0.003; Wilcoxon ranksum test. Significant comparisons shown in inset). All distributions were higher than their parallels during wakefulness (shown as dashed histograms; AUDp-U(3042,2915)=8.491e6 p<1e-4, TeA-U(529,196)=1.79e5 p<1e-4, AUDp-TeA-U(2296,918)=3.427e6 p<1e-4, Wilcoxon ranksum test). All distributions were different from shuffled (p<1e-4 for all comparisons, Kolmogorov-Smirnoff test). (h) Same as (g) for noise correlations (Median (IQR): AUDp-0.032 (0.005-0.052) n=2915, TeA-0.033 (0.007-0.071) n=198, AUDp-TeA – 0.022 (0.004-0.051) n=918). Correlations were highest in TeA (AUDp vs. TeA-U(2915,196)=4.502e6 p=0.006, AUDp vs. cross-region - U(2915,918)=5.598e6 p=0. 7 n.s, TeA vs. cross-region - U(196,918)=1.207e5 p=0.005; Wilcoxon ranksum test. Significant comparisons shown in inset), and generally larger than in the awake recordings (AUDp-U(3042,2915)=8.001e6 p<1e-4, TeA-U(529,196)=1.771e5 p<1e-4, AUDp-TeA-U(2296,918)=3.188e6 p<1e-4, Wilcoxon ranksum test). All distributions were different from shuffled (p<1e-4 for all comparisons, Wilcoxon ranksum test). (i) Scatter plot showing relationship between noise and signal correlations. Dashed lines show linear fits for each individual population. Linear correlation between the two was strongest for TeA and for cross region correlations (Linear fit, R^2^: AUDp-NC=0.14*SC+0.02 R^2^=0.257; TeA-NC=0.17*SC+0.02 R^2^=0.380; TeA-AUDp-NC=0.17*SC+0.02 R^2^=0.302).

The temporal aspects of the neural response were also changed in the anesthetized state. Latency was decreased in AUDp, although the hierarchical gradient was maintained (Fig. 7e; Minimal latency in TeA follows A1 by 11.5 ms, and latency to peak by 28.5 ms). Moreover, the temporal aspects of the late responses were particularly affected, such that the population response under anesthesia was shorter. AUDp and TeA retuned to baseline ~110ms and ~254 ms earlier as compared to the awake state, respectively (Fig. 7f). Suppressive population response were modulated as well, becoming weaker and generally more sluggish. However, the small number of SUs exhibiting suppressive responses under anesthesia in our dataset precluded a statistical validation of this qualitative impression (data not shown).

Next, we examined how anesthesia changes the interactions within and between neurons in AUDp and TeA. We expected that within-region correlations will be higher under anesthesia (Ecker et al. 2014; Harris and Thiele 2011). Indeed, both SCs and NCs were higher under anesthesia as compared to the awake state (Fig. 7g,h). While all correlations increased under anesthesia, the effect was larger in TeA and across-regions, making TeA SC and NC larger than their AUDp counterparts and cross-region correlations indistinguishable from AUDp correlations. Linear correlations between SC and NC increased as well (Fig. 7i). Together, the more prominent effects of anesthesia on TeA further suggests that TeA is a higher order auditory cortex.

## Discussion

The rodent’s TeA was recognized as an auditory field approximately thirty years ago. Initial anatomical studies of regions akin to TeA in the rat cortex showed feed-forward connectivity from primary auditory cortex, distinct connectivity patterns from the non-leminiscal auditory thalamus, and strong projections to the amygdala and striatum (LeDoux, Farb, and Romanski 1991; Romanski and LeDoux 1993; Arnault and Roger 1990). Recently, using trans-synaptic rabies tracing we found that TeA receives most of its inputs (~45%) from AUDp, as well as inputs from dorsal/medial thalamus that surpass those from ventral thalamus. In addition, TeA receives direct inputs from numerous other (>100) cortical and sub-cortical structures (Tasaka et al. 2020). Aside from describing its connectivity, physiological studies of TeA are scarce. Basic response profiles to pure tones have been documented in anesthetized rats with multi-unit recordings (Polley et al. 2007) and recently using surface imaging of the cortical sheet in mice (Romero et al. 2020). Here, using electrophysiology, we characterized how SUs in TeA represent pure tones and FMs in awake mice, and characterized interactions between TeA and its main upstream region, the primary auditory cortex. We suggest that TeA, by virtue of its physiological sparseness, temporal sluggishness, and improved representation of FMs is a high-order auditory processing station. Despite strong feed-forward connectivity evident from the anatomy (Tasaka et al. 2020), AUDp and TeA units exhibit weak functional correlations during wakefulness, suggesting TeA activity is strongly shaped by inputs from other regions, and/or within region computations.

### Extracellular recordings using Neuropixels probes

Neuropixels like other extracellular recording methods allow the collection of dozens of well isolated SUs. Exploiting the long shank of the Neuropixels probe (Jun et al. 2017), we were able to measure auditory responses along several stations of the cortical hierarchy, simultaneously. Notably, however, all methods have biases. Such biases must be recognized because they can be the reason for numerous discrepancies among studies. One such bias of extracellular recording methods is their tendency to capture specific cell types and particularly highly-active neurons (Barth and Poulet 2012). Indeed, comparing our data with previous reports where an unbiased recording method - like loose-patch recordings-was used, we found that both spontaneous firing rates and evoked responses tended to be larger for measurements with Neuropixels, in both awake and anesthetized preparations (Maor et al. 2020; L. Cohen and Mizrahi 2015; Hromadka T., R. DeWeese M. 2008). Another major source of variation is layer specificity. In our experiments, we penetrated the cortex to record from L5 in AUDp, and from various layers (albeit still dominated by L2/3 and L5) in TeA. Previous studies in rodents showed that AUDp L5 neurons are the least sparse along the cortical column, exhibiting broad tuning and short latencies competing and even preceding those of L4 neurons, which are classically considered the main recipients of lemniscal thalamic inputs (Sakata and Harris 2009; Wallace and Palmer 2008; Intskirveli et al. 2016). Studies in cat AUDp showed that feedback from auditory association areas is transmitted to all AUDp layers (Mitani and Shimokouchi 1985). Our own recent tracing experiments in mice showed that the input from AUDp to TeA is dominated by L5 (x3 times higher compared to L2/3), and feedback connections from TeA to AUDp were also dominated by L5-to-L5 projections (Tasaka et al. 2020). Thus, our recording setup is particularly prone to record from putatively interconnected neuronal populations in AUDp and TeA.

### Sparse activity in TeA

The most striking difference between the responses in TeA as compared to AUDp and AUDv was the increase in neural sparseness along the hierarchy. Both spontaneous and evoked activities were attenuated for all stimuli we presented, and sparseness increased in all the metrics we tested - i.e. bandwidth, population sparseness and lifetime sparseness (Fig. 2, Fig. 4, Fig. S5). Sparse coding is a guiding principle in neural coding. Theoretically, sparse coding has been suggested to have many advantages such as being energetically efficient, having a high memory and representational capacity, and facilitating efficient feature extraction from natural stimuli (Olshausen and Field 2004; Lewicki 2002; Laughlin and Sejnowski 2003). It has been widely reported that primary sensory cortices, auditory cortex included, are sparse in their nature (Hromadka T., R. DeWeese M. 2008; Barth and Poulet 2012; Vinje and Gallant 2000). While evidence from humans and primates showed that sparseness and responses to abstract features increase in higher-order sensory cortices (Quiroga et al. 2005; Waydo et al. 2006; Freiwald, Tsao, and Livingstone 2009; Wolfe, Houweling, and Brecht 2010), the few previous characterization of putative TeA actually showed that it was more broadly tuned as compared with AUDp (Polley et al. 2007; Romero et al. 2020). Our data from awake mice shows a different picture, and if anything, an opposite trend. We show that tuning in TeA narrows down and becomes more specific as compared to AUDp. Obvious differences between the studies which may underlie the observed differences are that we recorded in awake mice (see Fig. 7 for effects of anesthesia on sparseness and tuning bandwidth), and the recording methods. Here, we measured from SU while others measured from MUs or widefield imaging. In support of this claim, some metrics of sparseness that we tested on our MU data did not show statistically significant differences between AUDp and TeA (Table S3).

While the increase in sparseness along the cortical hierarchy is a well-documented phenomenon in other senses, the mechanisms underlying it are poorly understood. The most common suggested mechanisms argue that sparseness is a result of inhibition. For example, strong feed-forward or recurrent inhibition (Silberberg and Markram 2007; Kapfer et al. 2007), temporal dynamics between excitation and inhibition (Wehr and Zador 2003), and asymmetry between broadly-tuned inhibition and narrowly tuned excitation (Poo and Isaacson 2009; Haider, Häusser, and Carandini 2013), have all been argued to underlie sparser responses. In our data, suppressive responses were abundant in all cortical regions and in response to all presented stimuli (Fig. 2b; Fig. 4b), but not limited to TeA. Nevertheless, spontaneous and evoked activity were smaller in TeA (Fig. 2b,g), suggesting higher baseline levels of inhibition. While we do not fully understand how inhibition shapes responses in auditory cortex, it is clearly a prominent feature of the cortical response (Wehr and Zador 2003; Haider, Häusser, and Carandini 2013; Froemke 2015). Future characterization of inhibitory populations in TeA may provide a more mechanistic explanation to the source of sparseness in this brain region.

### TeA as a high-order auditory processing station

Previous work studying higher-order sensory cortices involved characterization of receptive fields, identification of neuronal feature selectivity profiles, and examination of activity during task engagement (Marshel et al. 2011; Andermann et al. 2011; Elgueda et al. 2019; Gilad, Maor, and Mizrahi 2020). Here, we studied auditory responses in passively listening mice, and identified two distinct features of FM responses in TeA - higher selectivity to FM direction (Fig. 5a) and decreased selectivity to FM speeds (Fig S8). The lower selectivity to FM speed is possibly in agreement with previous reports of invariance to spectro-temporal modulations of sounds in the rat SRAF (presumably the equivalent of mouse TeA; Carruthers et al. 2015b). Alongside the SU response features to FMs, population discriminability for FM stimuli was larger in TeA compared to AUDp, and opposite to the trend observed for pure tones. Increased selectivity and discriminability for complex sensory stimuli is a prominent feature of higher-order visual cortices (Marshel et al. 2011; Freiwald, Tsao, and Livingstone 2009), supporting a similar role for TeA in audition.

Classically, longer response latencies are associated with higher-order processing (Bullier and Nowak 1995; Capalbo, Postma, and Goebel 2008), but latencies were also suggested to underlie a neural latency code (Paoli et al. 2018; Shriki, Kohn, and Shamir 2012; Storchi et al. 2012) or even be correlated with the degree of conscious perception (Reber et al. 2017) and stimulus selectivity (Mormann et al. 2008). Since TeA receives the vast majority of its inputs directly from AUDp, and particularly from L5, we expected an order of ~15 ms difference in response latency between the two regions (London et al. 2010). Indeed, the minimal latencies of SUs in both regions showed just that (Δmedians=14 ms; Fig. 3b). Aside from minimal latency, the ‘time to peak’ of responses and the overall response duration are larger in TeA (Fig. 3c,f). This temporal profile of responses is also compatible with TeA being a higher-order auditory processing station, directly downstream of AUDp (Elgueda et al. 2019).

To the best of our knowledge, only two studies causally manipulated TeA to test its function in behaving animals. TeA was shown to be necessary for auditory fear conditioning (Dalmay et al. 2019), and for maternal recognition of pup calls (Tasaka et al. 2020). Both these functions of TeA may supervene, at least partially, on strong bi-lateral projections between TeA and the amygdala (Tasaka et al. 2020; Dalmay et al. 2019; Tsukano et al. 2019; LeDoux, Farb, and Romanski 1991). We postulate that TeA’s unique connectivity profile (Tasaka et al. 2020), and physiological signature as a high-order station, enables it to take a more central part in computations that integrate auditory information with other cues, which may be experience dependent. Characterization of TeA’s activity during auditory-task performance and its manipulation in additional behavioral contexts and following learning will shed further light on its functional role and the specific computations it contributes to.

### Functional connectivity between AUDp and TeA

Ketamine-Domitor anesthesia affected both AUDp and TeA. However, TeA’s activity was significantly more modulated by anesthesia. Specifically, activity in TeA increased, sparseness decreased, and auditory responses were faster and lasted for shorter durations (Fig. 7). Ketamine anesthesia is known to leave primary sensory representations intact, affecting mostly activity in frontal regions and top-down functional connectivity (Mashour 2014; Blain-Moraes et al. 2014; Schroeder et al. 2016). One of the prominent changes induced by anesthesia is the decrease in late post-stimulus activity and shortening of the auditory response (Fig. 7f). Accumulating evidence show that late sensory-evoked activity is associated with conscious perception of stimuli and behavioral outputs (Sachidhanandam et al. 2013; Del-Cul, Baillet, and Dehaene 2007). In addition, it was demonstrated that late post-stimulus dendritic spikes and bursting activity in L5 pyramidal cells in somatosensory cortex are induced via top-down excitatory connections targeting the apical tuft (Manita et al. 2015). We, therefore, suggest that a decrease in top-down inputs is the main cause of response shortening in auditory cortex. This hypothesis is in agreement with reports showing that late but not early activity is abolished by general anesthesia (Hudetz, Vizuete, and Imas 2009). Thus, anesthesia likely shuts down much of the rich input landscape to TeA leaving only early feed-forward signals intact.

## Methods

### Extracellular recordings using Neuropixels

10-13-week-old *TRAP2;TB* double heterozygous females (F1 hybrid of C57BL/6 and FVB strain; n=5 awake mice, n=5 anesthetized mice) were used for electrophysiological recordings. Prior to the recording, a custom-made metal bar was implanted on the mouse skull, and a small craniotomy was made on the left hemisphere (coordinates relative to Bregma: anterior −2.5, lateral 4.2 mm). The craniotomy was protected by a wall of dental cement and covered with a silicone elastomer (WPI; Kwik-Cast cat#KWIK-CAST). The surgery was done under Ketamine Medetomidine anesthesia (0.80 and 0.65 mg/kg respectively) and a subcutaneous injection of Carprofen (0.004 mg/g). For awake recordings, mice were given 1-2 days to recover, after which they were head-fixed for about 30 min to habituate to the recording setup. Recordings were performed 1-2 days post habituation. On the day of the recording, animals were head-fixed and the craniotomy was exposed. Then, a Neuropixels probe (imec, phase 3A) was inserted through the craniotomy in a 20-degree tilt (from vertical position) and lowered down approximately 3 mm deep. Penetration and probe depth were performed and monitored using a micromanipulator (Scientifica PatchStar micromanipulator). Probes were covered with a fluorescent dye (DiI [Invitrogen cat#V22885] or DiO [Invitrogen cat#V22886]) before penetration, to enable reconstruction of penetration sites in high resolution. For anesthetized recordings, the procedure was the same but followed the surgery directly, under continuous anesthesia. In each mouse, we performed up to three consecutive probe penetrations. Probe trajectories were reconstructed from consecutive coronal slices using an open source software (Shamash et al. 2018; Fig 1d, Fig S1). Following the reconstruction, probe channels were annotated with corresponding brain regions they were recording from.

All recordings were acquired using Neuropixels phase 3A probes (imec), a base-station connector (imec) and a commercially available FPGA board (KC705, Xilinx). An external reference electrode (Ag/AgCl wire) was positioned on the skull and covered with saline solution. Data was sampled at 30 kHz, with action potential band filtered to contain 0.3-10 kHz frequencies. Action potential band gain was set to 500. All recordings from the same animal and position were concatenated, and automatically sorted using Kilosort/Kilosort2 (anesthetized and awake correspondingly) open source software (Pachitariu et al. 2016; https://github.com/MouseLand/Kilosort2). Following automatic sorting, manual sorting was performed using ‘Phy’ open-source GUI (UCL; https://github.com/cortex-lab/phy). During manual sorting, spike clusters were merged based on assessment of wave-form similarity and the appearance of drift patterns. Finally, each spike cluster was assessed in criteria of waveform size, consistency, and the presence of short-latency inter-spike-intervals (ISIs). If and only if a cluster was satisfactory on all accounts, it was tagged as a single unit (SU) corresponding to a single neuron. If a cluster did not meet the above criteria but was clearly reflecting neuronal activity (and not noise) it was tagged as a multi unit (MU), corresponding to a group of neurons clustered together. Figures 1d and S1 were generated using the Allen CCF software (UCL;https://github.com/cortex-lab/allenCCF). Figure 1e was generated using the Neuropixels-utils software kit, developed by Dan O’shea (https://github.com/dioshea/neuropixel-utils).

### Auditory stimuli

The pure tone protocol was comprised of 30 pure tones (100 ms duration, 5 ms on and off linear ramps) logarithmically spaced between 3 and 80 kHz, presented at four sound pressure levels (72-42 dB SPLs). Each frequency x attenuation combination was presented 12 times. Pure tones were presented in a random order, with 600 ms intervals between tone onsets.

The FM protocol was comprised of 20 stimuli, spanning the range between 4-64 kHz either in a positive (“rising”) of negative (“falling”) frequency modulation (1 ms on and off linear ramps). 10 stimuli had linear frequency modulation, with varying slopes and corresponding stimulus times (±30 kHz/sec, 2 sec; ±79.78 kHz/sec 752 ms; ±212.16 kHz/sec 283 ms; ±563.91 kHz/sec 106 ms; ±1500 kHz/sec 40 ms). The other 10 stimuli had logarithmic slopes that generated stimuli with matching durations (±2 Oct/sec, ±5.32 Oct/sec, ±14.14 Oct/sec, ±37.61 Oct/sec, ±100 Oct/sec). All stimuli were presented in 62 dB SPL for 12 repetitions. Stimuli were presented at 0.37 Hz in randomized order.

### Pure tone data analysis

Data analysis and statistics were performed using a custom written code in MATLAB (MathWorks). Prior to analysis, multi units and single units firing less than 10 spikes during the entire protocol were eliminated. Spontaneous firing rates were calculated from baseline activity (100 ms) preceding all trials (n=1440). To detect auditory units, we looked for 50 ms maximal and minimal response windows in the time between 0 to 200 ms post stimulus onset. The maximal and minimal response windows were then compared to 50 ms of baseline (one-sided two-sample t-test, p<0.05). According to test results, units were considered as excited and or suppressed. If no significant window was found, the unit was considered non-auditory and excluded from further analysis. Based on these response windows, frequency-response areas were extracted (FRAs), and the best-frequency (BF) of excited units was identified as the frequency evoking the maximal response (in a single attenuation). Evoked firing rates (FRs) were considered as FRs in response to a units’ BF during the maximal response window, subtracted with the units’ spontaneous FR.

To determine the significance of responses to individual frequency x attenuation combinations, response during the optimal (excited/suppressed) windows was compared to baseline activity in the relevant trials via one-tailed Wilcoxon ranksum tests. Bandwidth of excited units was calculated as the number of frequency x attenuation combinations eliciting a significant response, subtracted with the number of expected false positives based on the number of performed comparisons (30 frequencies. x 4 attenuations = 120) and the required significance level(p<0.05). Population sparseness was then calculated as the fraction of units in each region exhibiting a significant response to each one of the stimuli. Lifetime sparseness was calculated for excited units from firing rates to all tones in a given attenuation (chosen to be the attenuation of the BF) using the following formula:

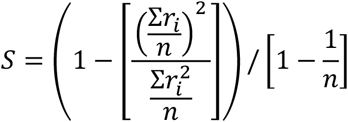

The values *r_i_* are the firing rates in response to each frequency, and n=30 is the number of pure tones presented. Values of S near 0 correspond to a dense code and values near 1 correspond to a sparse code (Vinje and Gallant 2000).

Latency to peak was calculated as the time post-stimulus in which the units’ PSTH (smoothed with a 9 ms window) is maximal. A similar approach was applied for time to minima of suppressed units. To calculate minimal latency, we first screened for units having a significant early excitatory response window, by looking for a significant 50 ms excitatory response window within 0-110 ms post stimulus onset (similarly to what was described above). We calculated FRAs for the early response windows, and extracted the “early BF”. The trials corresponding to the early BF were isolated (only trials from the best attenuation), and the latency of the first post-stimulus spike in each trial was detected. The minimal latency of a unit was considered the early BFs’ median latency to first spike.

Population response was calculated by averaging the trials of all units in a given region to each one of the pure tone stimuli (frequency x attenuation combination). After averaging, the mean baseline response (100 ms pre stimulus onset) was subtracted from the trace, and the response was smoothed (9 ms window). Finally, the population responses to all stimuli were averaged together to obtain mean and SEM of the population response to pure tones. To assess deviation from baseline, the population response at every given 1 ms time bin was compared via a two-sample t-test to the mean baseline population response (average of the activity during 100 ms pre stimulus). Required significance level was adjusted to multiple comparisons by requiring p<0.05/#compared time bins (0.05/700).

### FM analysis

Here too, data analysis and statistics were performed using a custom written code in MATLAB (MathWorks). Prior to analysis, multi units and single units firing less than 10 spikes during the entire protocol were eliminated. First, and similarly to pure tones, maximal and minimal 50 ms response windows were detected in response to each one of the FM stimuli. For each such window, the firing rate within the window was calculated, and the response was compared to 50 msec of baseline in a one-tailed Wilcoxon rank sum test. This determined whether a unit was significantly excited/suppressed by each FM stimulus (total of 20 stimuli). Units were considered responsive to FMs if at least one FM stimulus elicited a significant excitatory/suppressive response. A unit was considered excited if it had a significant excitatory window to at least one FM stimulus, and suppressed if the same was true for suppressive windows. If both types of windows were significant for at least one stimulus, the unit was considered both excited and suppressed. FM bandwidth was considered the number of FM stimuli (out of total 20) eliciting a significant excitatory response. Population sparseness and lifetime sparseness were calculated similarly to pure tones based on responses to all FM stimuli.

Direction selectivity index (DSI) was calculating using 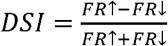, where *FR* ↟ refers to the summed firing rate for all rising FMs, and *FR* ↓ is the summed firing rate for all falling FMs. Firing rates were calculated based on maximal 50 ms response window to each FM stimulus. Classification ability to rising vs. falling FMs was estimated using two classifiers – SVM (with a linear Kernel) and 5-nearest neighbors. For both classifiers, 5000 iterations of training and test were conducted to assess classification performance. In each iteration, all trials from n=30 randomly selected units from each region were pooled and divided with 5:1 ratio to train and test trials. Performance was assessed by monitoring the test error rate over all iterations. To assess classifiers success under matching firing rate conditions we ran the classifiers again with a constraint on the neurons being pooled for each train-test iteration. We calculated the firing rate distribution of TeA units, and divided it to three quantiles. In each iteration of the classifier, we validated that for each region (AUDp, AUDv and TeA) a third of the used units have firing rates matching TeA’s lowest quantile, a third matches the center quantile and a third matches the top quantile. The constraint was applied also on TeA units. Practically, this led to higher sampling of weakly activated units in all regions. To calculate population speed selectivity, each unit’s response to FMs with same slope and different directions was summed (responses calculated based on maximal response windows for each FM as mentioned before). All units within each region were pooled, and the population response to different FM speeds was pairwise compared via a two-sampled t-test. P-value significance level was Bonferroni corrected to the number of performed comparisons (0.05/10).

FM Minimal latency and latency to peak were calculated similarly to pure tones, only using FM stimuli trials and PSTHs.

### Correlation analysis

Pairwise signal correlations were calculated by computing the Pearson correlation of FRAs between every couple of simultaneously recorded units. Note that since FRAs were calculated based on response windows unique to each unit, correlated variability (noise correlation) should not contribute to the calculated signal correlations. FRAs were constructed based on excitatory response windows for units having them and based on suppressive response windows for units lacking excitatory responses. Shuffled distributions were calculated by correlating shuffled FRAs, and repeating this for n=500 iterations. To calculate noise correlations, data was binned into 20 ms bins. For each pure tone stimulus (frequency x attenuation combination) the mean response in all time bins was calculated, and subtracted from all individual trials to obtain “trial-fluctuation-vectors”. For each unit, fluctuations from 0-200 ms (10 time bins) post stimulus (from all stimulus trials) were concatenated to form a long fluctuation vector. Pairwise noise correlations were calculated via Pearson correlating between the concatenated fluctuation vectors. Shuffled noise correlations were calculated by randomizing the time bins in each units’ fluctuation vector and calculate the resulting Pearson correlation. This procedure was repeated for n=500 iterations.

### d prime analysis

In order to estimate the ability of a network to discriminate between two given pure tones or FM sweeps, we calculated pairwise d primes based on the activity of the entire SU population recorded in each region. For two given auditory stimuli (either two pure tones or two FM sweeps), p and q, d primes were calculated according to the formula:

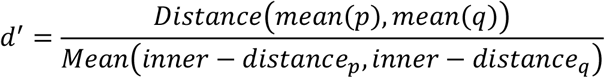

where p/q is a matrix with 12 columns (one column for each repetition) and n rows (n = number of SUs in each cortical region), in which each entry contains the response of a specific SU to a specific stimulus at a specific trial. mean(p)/mean(q) are vectors with n entries representing the mean coordinates (averaged over the individual trials) in n-dimensional space for stimuli p/q. The difference between these vectors was assessed as a scalar by Euclidean distance, i.e. 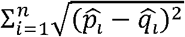. Inner-distance_p and inner-distance_q were similarly assessed as two scalars, by calculating the Euclidean distance between each one of the 12 single trials with the stimulus average representation, and then averaging over the 12 resulting measures.

### Statistical analysis

All statistical analyses were performed with MATLAB (MathWorks). Across-region comparisons were conducted using Kruskal-Wallis test, followed up with post hoc Tuckey-Davies Honest Significant Test unless mentioned otherwise. Testing distributions for normality was performed using Lilliefors test. Comparisons of distribution “shapes” were usually performed using Kolmogorov-Smirnoff tests.

## Code Accessibility

Codes used for data analysis are available from the corresponding author upon request or in https://github.com/LibiF/Neuropixels-Scripts.

## Acknowledgements

We thank members of the Mizrahi laboratory for comments on the manuscript. We thank the Gatsby Foundation for partnering in the development of neuropixels and providing access to engineering prototype probes. We thank HHMI Janelia and UCL for helpful discussion, the development of data acquisition and analysis tools, and training in the use of Neuropixels. We gratefully acknowledge the support of the NVIDIA Corporation with the donation of the Titan Xp graphics processing unit (GPU) used for this research and the analysis of Neuropixels data. This work was supported by an ERC consolidators grant to A.M. (#616063), Israeli Science Foundation grant to A.M. (#224/17), and the Gatsby Charitable Foundation.

## Author contributions

L.F. and A.M. designed the experiments. L.F., G.T., and I.M. conducted the experiments. L.F. analyzed the data. L.F and A.M. wrote the paper.

## Supplementary Figure legends

**Figure S1 - Reconstructed probe tracks**

Reconstructed anatomical tracts of 12 probe penetrations used to record the electrophysiological data from awake mice. The Y axis describes depth and corresponding brain region. The X axis describes the distance to the nearest brain region (larger distance implies greater separation between regions). We analyzed data from three brain regions only - AUDp (blue), AUDv (purple) and TEa (green). Data was collected from 5 awake head-fixed mice (mouse 1: probes 1-3; mouse 2: probes 4-6; mouse 3: probes 7-8; mouse 4: probes 9-10; mouse 5: probes 11-12). Pure tone data was collected from all mice, while FM data was collected from mice 2-5. Acronyms: VISrl-Rostrolateral visual area; VISal-Anterolateral visual area; AUDd-Dorsal auditory area; AUDpo-Posterior auditory area; AUDp-Primary auditory area; AUDv-Ventral auditory area; TEa-Temporal association areas; ECT-Ectorhinal area; PERI-Perirhinal area; SSp-bfd-Primary somatosensory area, barrel field structure; SSs-Supplemental somatosensory area. Numbers refer to the cortical layer within the given structures.

**Figure S2 - Spontaneous firing rates are distributed log-normally**

Spontaneous firing rate distributions in auditory cortices in linear (top) and logarithmic (bottom) scale (Median (IQR): AUDp – 1.13 (0.21-3.48) Hz n=240, AUDv-0.56 (0.14-1.63) Hz n=218, TeA– 0.19 (0.07-0.67) Hz n=97). Spontaneous FRs of SUs in all brain regions were log-normally distributed (AUDp-K(240)=0.058, p=0.03 n.s; AUDv-K(218)=0.04, p=0.5 n.s; TeA-K(97)=0.073, p=0.23 n.s; Lilliefors test for normality Bonferroni corrected for MC).

**Figure S3 - Diversity of pure tone responses**

(a)-(i) Examples of SU responses to pure tones. Raster plots describe responses to 30 one hundred ms pure tones logarithmically spaced between 3-80 kHz, at 4 sound attenuations (10-40 dB attenuation). Upper axis shows the unit PSTH (scale is adjusted on a per SU basis). Stimulus is presented at time 0-100 ms (gray shaded region), and each units’ maximal (red) or minimal (blue) fifty ms response window is marked. Unit (a) was recorded in AUDp, and all other units recorded in TeA.

**Figure S4 - Distributions of BFs and evoked responses**

(a) SU best frequency (BF) distribution by recorded region and probe penetration. Each point represents the BF of one SU, with colors corresponding to unique probe penetrations (units sharing color were simultaneously recorded). BF distribution in all regions do not vary across penetrations (H(11)=18.92, p=0.06; Kruskal-Wallis), however within region one AUDv recording (#7) was biased towards high BF ranges (AUDp-H(11)=9.97, p=0.53 n.s; AUDv-H(11)=32.74, p=0.0006; TeA-H(11)=10.46, p=0.23 n.s; Kruskal-Wallis). (b) BFs difference in octaves of adjacent SUs (≤8 contacts apart or up to ~80 *μm* distance) originating from the same or different (“Within region”/Across regions”) brain regions (Median (IQR): Within region-1.31 (0.49-2.29) octaves, n=880 pairs; Across regions – 2.12 (0.49-2.87), n=69 pairs). Transition between auditory cortices induces increased shifts in units’ BFs (U=4.110e5, p=0.0013; Wilcoxon rank sum test). (c) Evoked firing rate distributions are distributed Log-normally. Evoked responses at the BFs are shown in linear (top) and logarithmic (bottom) scales (Median (IQR): AUDp-24.51 (13.89-43.30) Hz n=164, AUDv-21.24 (13.07-41.67) Hz n=136, TeA-11.44 (8.17-37.17) Hz n=55). FRs were log-normally distributed in all regions (AUDp-K(164)=0.073, p=0.03 n.s; AUDv-K(136)=0.057, p=0.33 n.s; TeA-K(55)=0.127, p=0.03 n.s; Lilliefors test for normality Bonferroni corrected for MC).

**Figure S5 - Development of sparseness from AUDp to TeA**

(a) Heat-maps showing the fraction of significantly responsive SUs to all presented pure tones (4 attenuations x 30 frequencies) in AUDp (top), AUDv (center) and TeA (bottom). Color-bar applies to all three heat-maps. (b) Boxplot summarizing the distribution of SU response probabilities shown in ‘a’ (Median (IQR): AUDp-0.16 (0.10-0.20), AUDv-0.13 (0.06-0.18), TeA-0.08 (0.06-0.12)). Response probability is highest in AUDp and lowest in TeA (H(2)=40.85 p<1e-4; Kruskal-Wallis. AUDp vs. AUDv-0.02, AUDp vs. TeA-p<1e-4, AUDv vs. TeA-p=0.007; TK HSD test). (c) Scatter plot showing relationship between SU latency to peak and it’s lifetime sparseness (LS). Individual units are color-coded according to anatomical affiliation (blue-AUDp, purple-AUDv, green-TeA). Dashed black line shows calculated linear regression (lat.=LS*74.90+57.89 R2=0.046). The two parameters were significantly correlated (r=0.207, p<1e-4).

**Figure S6 - Responses to linear and logarithmic FM are not different**

𝛥Number of significant stimuli per SU (lin. FMs - log. FMs), median (IQR): AUDp - 0 (−1-1) n =186, AUDv - 0 (−1-1) n=174, AUDp - 0 (0-1) n=70). For all regions, delta’s are centered around zero (AUDp-p=0.18 n.s, AUDv-p=0.45 n.s, TeA-p=0.55 n.s; Wilcoxon signed rank test), and distributions are indifferent from one another (H(2)=1.67 p=0.43 n.s; Kruskal-Wallis).

**Figure S7 - Diversity of FM responses**

(a)-(l) Examples of SU responses to linear (left column) and logarithmic (right column) FMs. Shaded regions mark FM stimuli durations. Y axis labels correspond to FM slopes in kHz/s(left) or Oct/s (right). Examples (a)-(d) SUs recorded in AUDp, (e)-(h) SUs recorded in AUDv and, (i)-(l) SUs recorded in TeA.

**Figure S8 - Speed selectivity for logarithmic FMs**

(a) Normalized population FRs for different logarithmic FM slopes (AUDp-n=162 SUs, AUDv-n=132 SUs, TeA-n=35 SUs). Bold lines and shaded regions mark mean response and SEM, respectively (blue-AUDp, purple-AUDv, green-TeA). Insets show statistical comparisons of within population responses to different FM speeds. (b) Tables showing p-values corresponding to tables shown a. P-values calculated based on two-sample t-tests, with significance level Bonferroni corrected for multiple comparisons by setting p<0.005.

