## Supplemental figures and Tables for "Sparse Coding in Temporal Association Cortex Improves Complex Sound Discriminability"

### Figure S1 - Reconstructed probe tracks

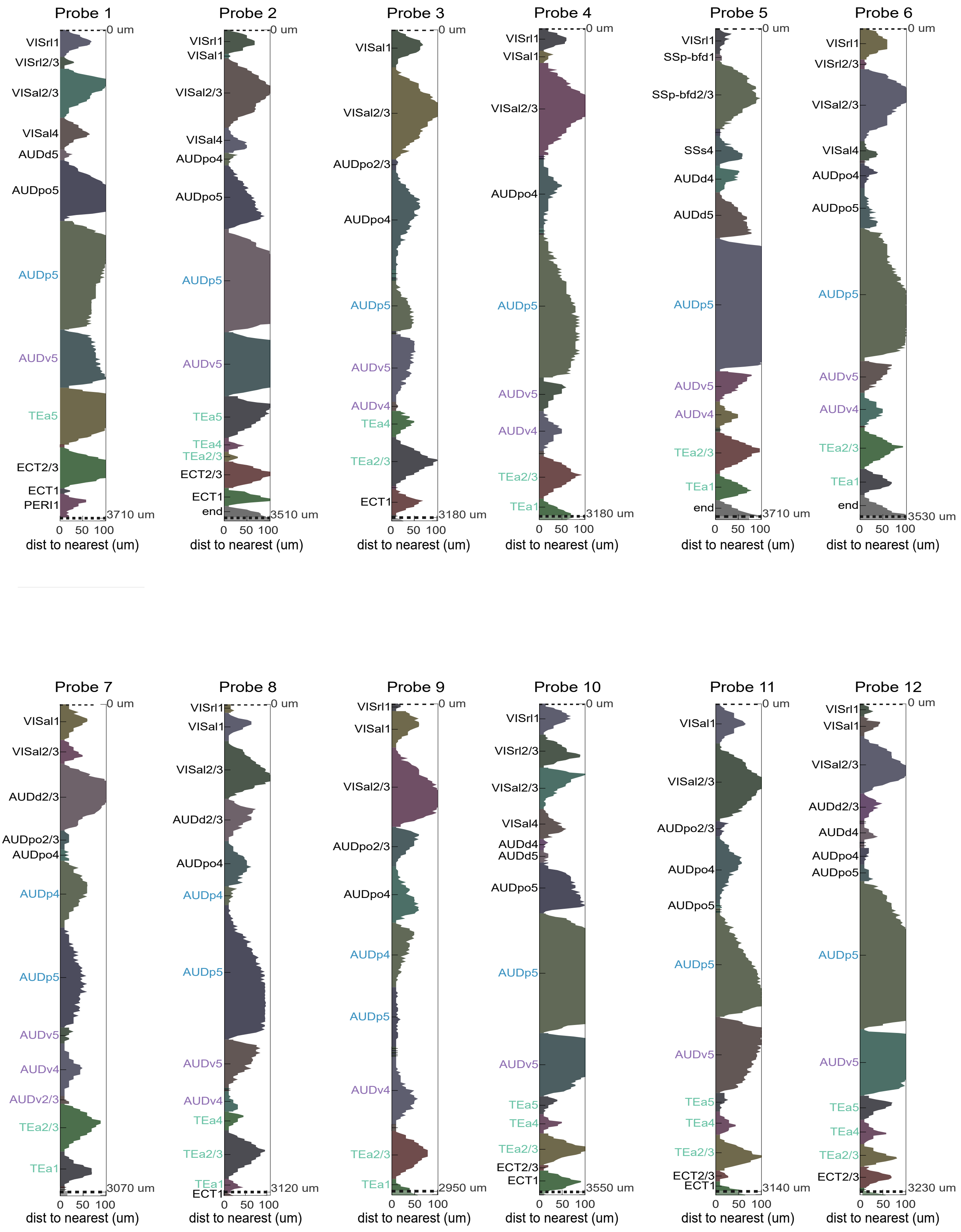

### Figure S2 - Spontaneous firing rates are distributed log-normally

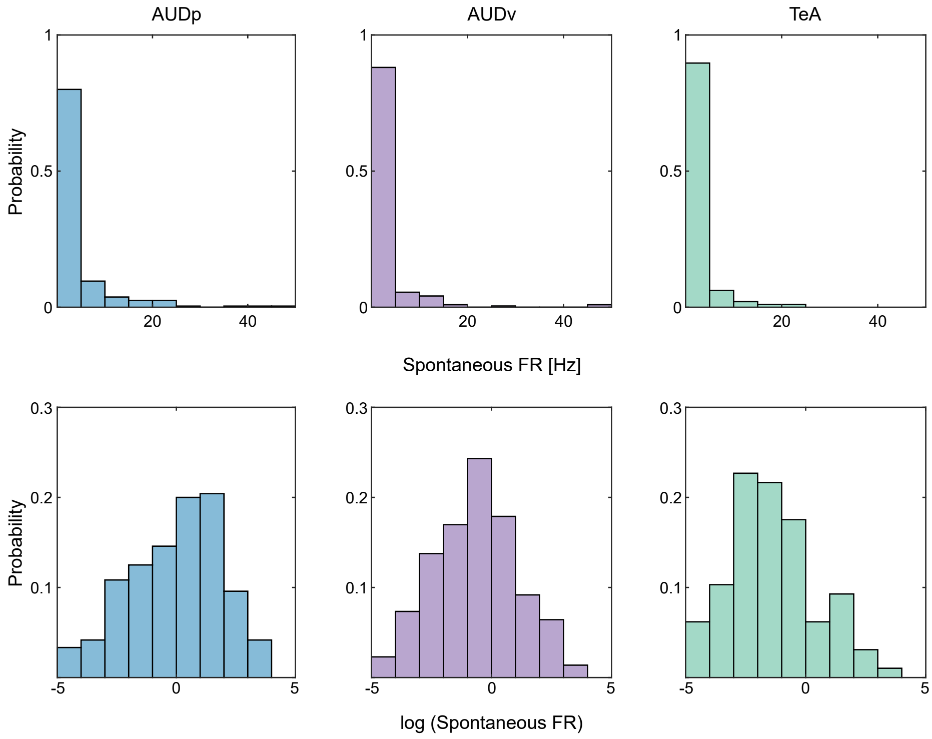

### Figure S3 - Diversity of pure tone responses

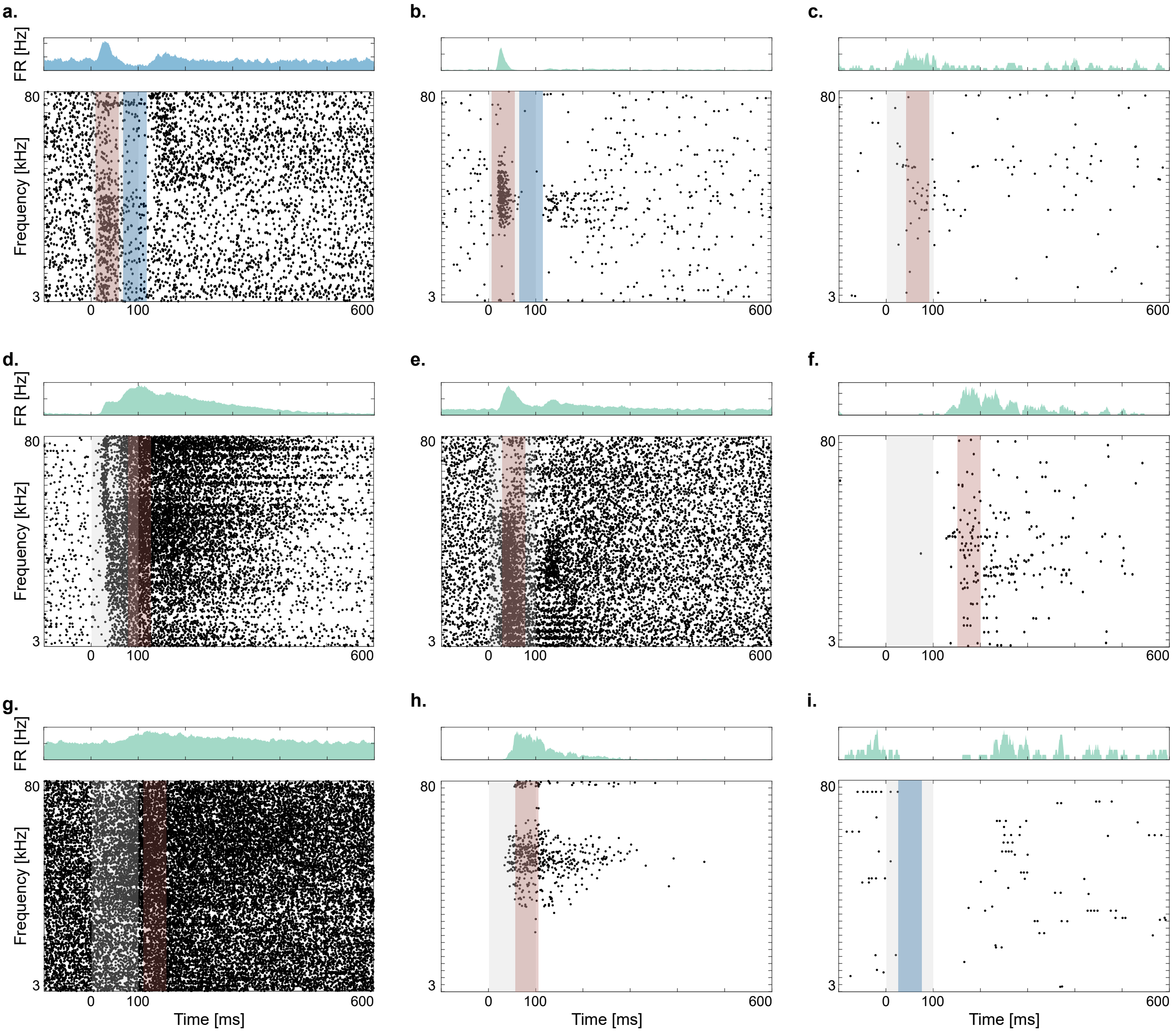

### Figure S4 - Distribution of BFs and evoked responses

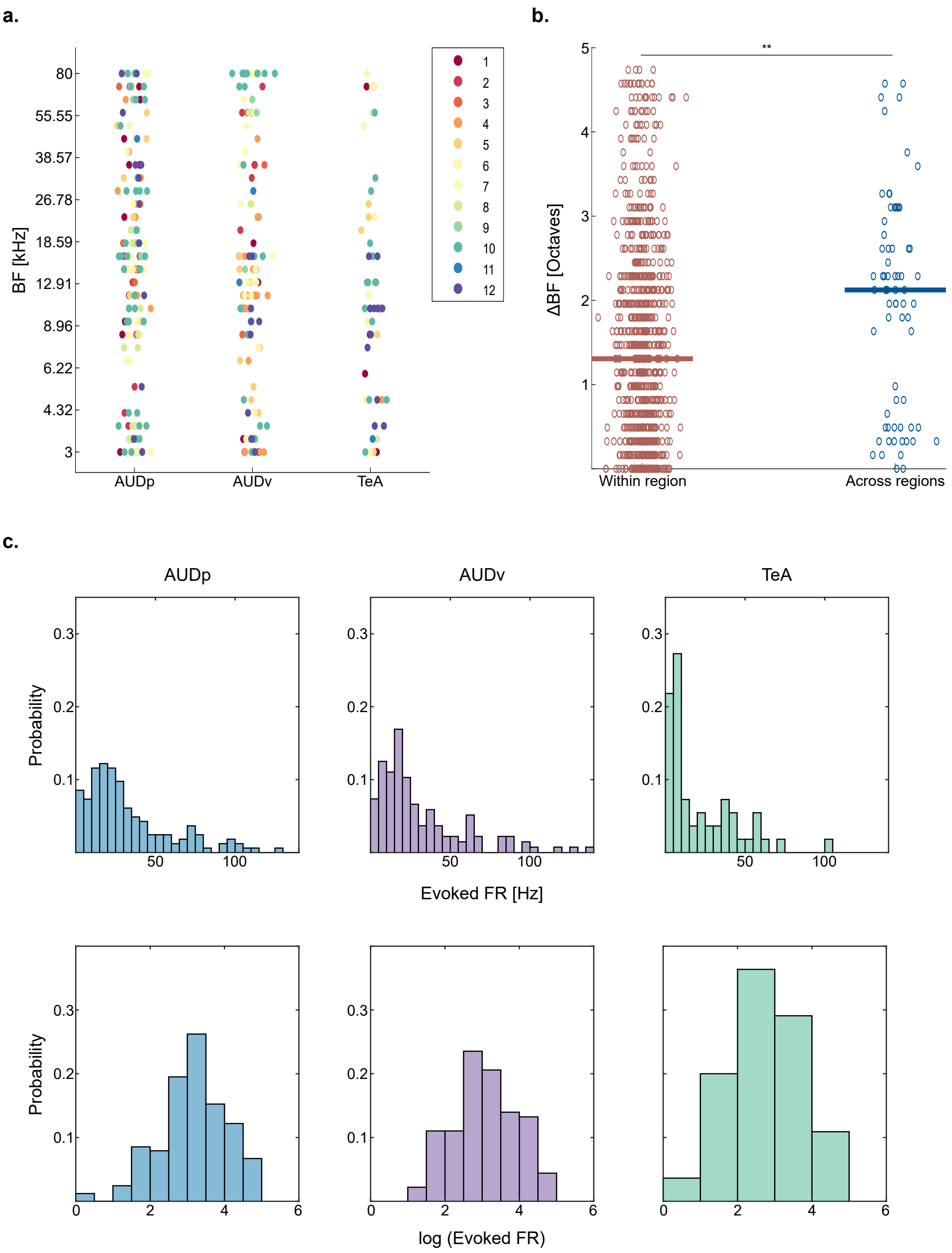

### Figure S5 - Development of sparseness from AUDp to TeA

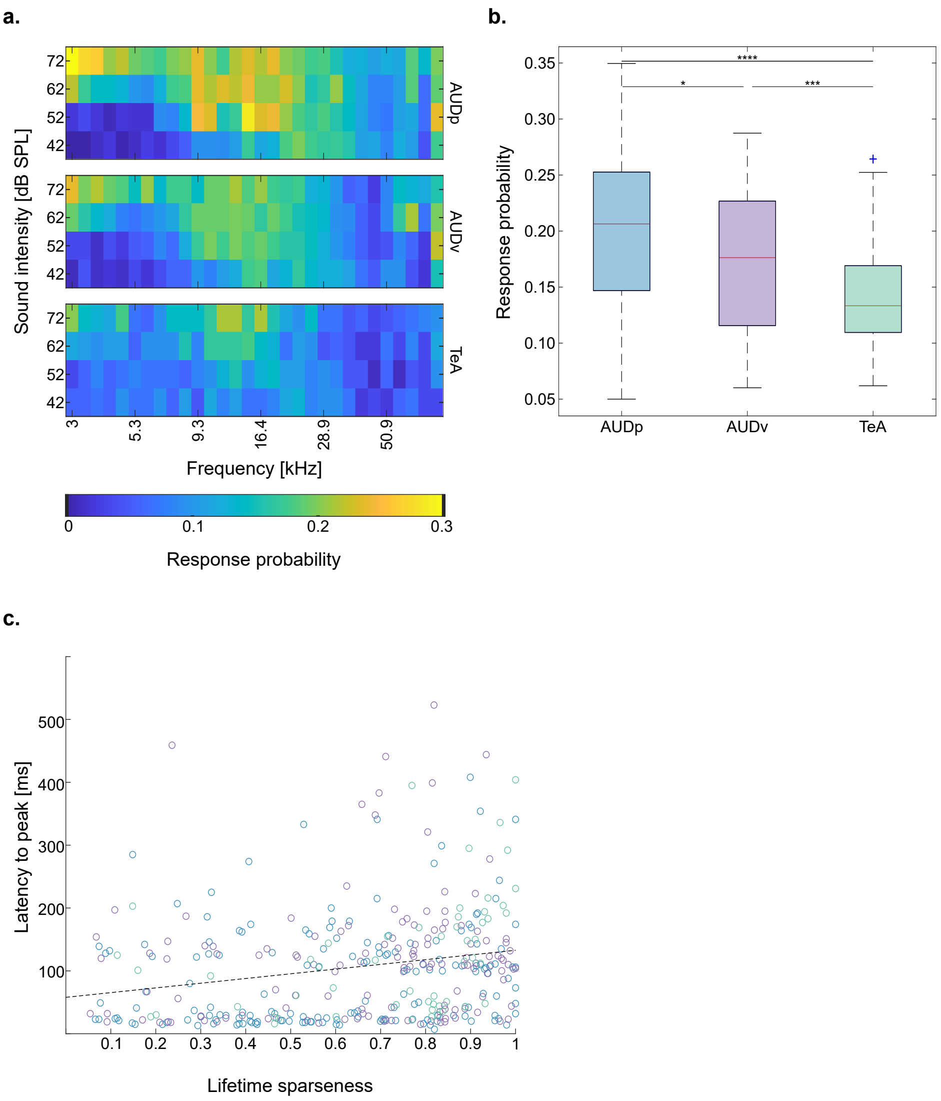

### Figure S6 - Responses to linear and logarithmic FMs are not different

**a.**

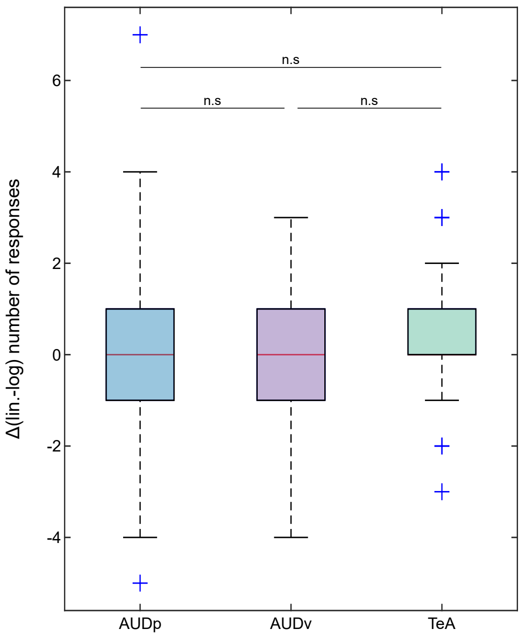

### Figure S7 - Diversity of FM responses

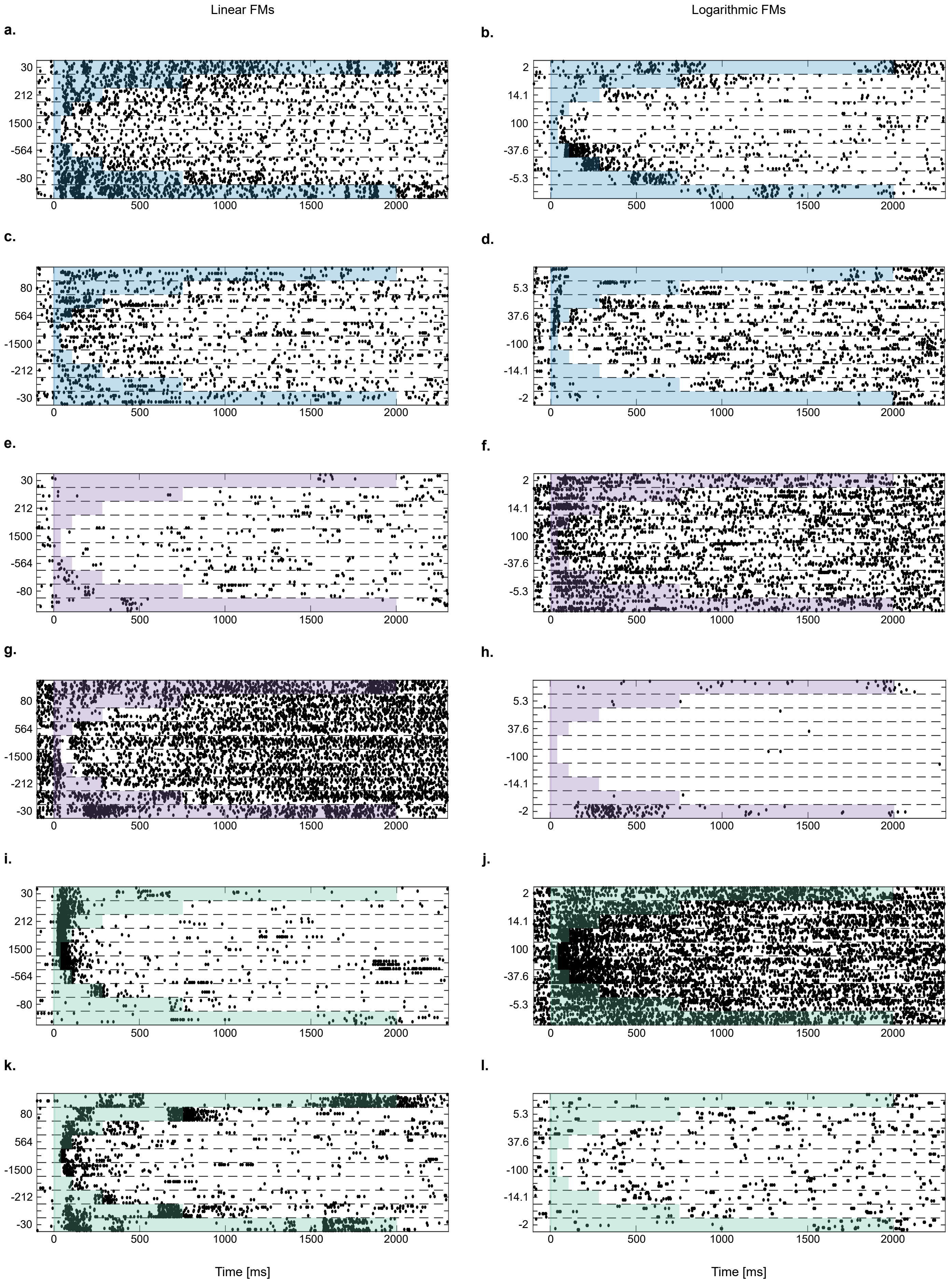

### Figure S8 - Speed selectivity for logarithmic FMs

**a.**

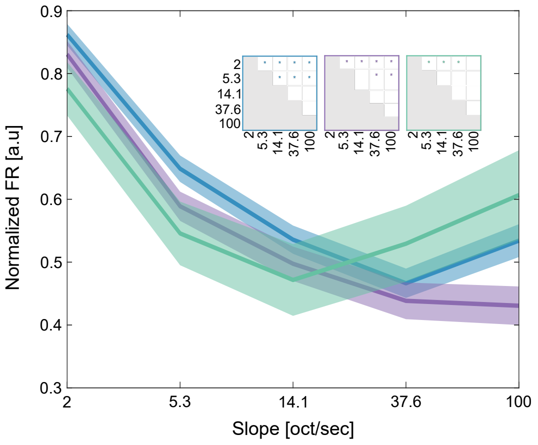


**b.**

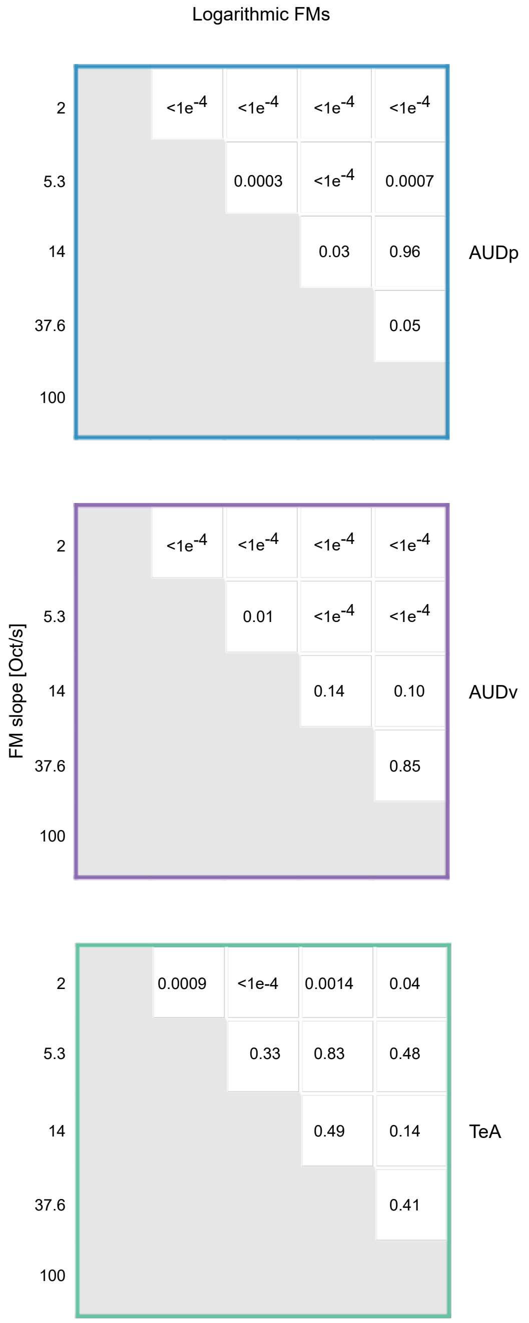

#### Table 1 – SU recording statistics

| | Pure Tone protocol | | FM protocol | | | |
|---|---|---|---|---|---|---|
| | #Num recorded units | Responding to PTs | #Num recorded units | Responding to linear FMs | Responding to logarithmic FMs | Responding to any FM |
| **AUDp** | 240 | 227 (95%) | 186 | 167 (90%) | 164 (88%) | 174 (93%) |
| **AUDv** | 218 | 198 (91%) | 174 | 137(79%) | 135(78%) | 142 (82%) |
| **TeA** | 97 | 84 (87%) | 70 | 42 (60%) | 36 (51%) | 45 (64%) |

### Table 2 – SU pure tone response statistics

| | Spont. FR [Hz] | Bandwidth [#num] | Lifetime sparseness | BF [kHz] | Evoked FR [Hz] | Minimal latency [msec] | Latency to peak [msec] |
|---|---|---|---|---|---|---|---|
| **AUDp** | 1.1 (0.2-3.5) | 11 (0-30) | 0.63 (0.39-0.83) | 15.5 (8.3-32.3) | 24.5 (13.9-43.3) | 34 (21-59) | 76 (23-135) |
| **AUDv** | 0.6 (0.1-1.6) | 4 (0-28) | 0.75 (0.51-0.87) | 13.8 (7.4-53.9) | 21.2 (13.1-41.7) | 48 (26-76) | 110 (34-149) |
| **TeA** | 0.2 (0.1-0.7) | 0 (0-17) | 0.82 (0.59-0.90) | 10.4 (6.3-20.0) | 11.4 (8.2-37.2) | 48 (33-73) | 112 (43-181) |

### Table S1 – MU recording statistics

| | Pure Tone protocol | | FM protocol | | | |
|---|---|---|---|---|---|---|
| | #Num recorded units | Responding to PTs | #Num recorded units | Responding to linear FMs | Responding to logarithmic FMs | Responding to any FM |
| **AUDp** | 190 | 185 (97%) | 158 | 147 (93%) | 151 (96%) | 153 (97%) |
| **AUDv** | 165 | 152 (92%) | 128 | 109 (85%) | 107 (84%) | 116 (91%) |
| **TeA** | 96 | 87 (91%) | 80 | 50 (62%) | 55 (69%) | 56 (70%) |

### Table S2 – Unit layer distributions

| | | Pure Tone protocol | | | | FM protocol | | | |
|---|---|---|---|---|---|---|---|---|---|
| | Unit type | #Num recorded units | Units per layer | | Percent of total units [%] | #Num recorded units | Units per layer | | Percent of total units [%] |
| **AUDp** | SU | 240 | 1 | - | - | 186 | 1 | - | - |
| | | | 2/3 | - | - | | 2/3 | - | - |
| | | | 4 | 2 | 0.8 | | 4 | 2 | 1 |
| | | | 5 | 238 | 99.2 | | 5 | 184 | 99 |
| | | | 6 | - | - | | 6 | - | - |
| | MU | 190 | 1 | - | - | 158 | 1 | - | - |
| | | | 2/3 | - | - | | 2/3 | - | - |
| | | | 4 | 2 | 1 | | 4 | 3 | 2 |
| | | | 5 | 188 | 99 | | 5 | 155 | 98 |
| | | | 6 | - | - | | 6 | - | - |
| **AUDv** | SU | 218 | 1 | - | - | 174 | 1 | - | - |
| | | | 2/3 | 2 | 0.9 | | 2/3 | 2 | 1.1 |
| | | | 4 | 69 | 31.7 | | 4 | 69 | 39.7 |
| | | | 5 | 147 | 67.4 | | 5 | 103 | 59.2 |
| | | | 6 | - | - | | 6 | - | - |
| | MU | 165 | 1 | - | - | 128 | 1 | - | - |
| | | | 2/3 | 8 | 4.8 | | 2/3 | 8 | 6.2 |
| | | | 4 | 39 | 23.6 | | 4 | 37 | 28.9 |
| | | | 5 | 118 | 71.5 | | 5 | 83 | 64.8 |
| | | | 6 | - | - | | 6 | - | - |
| **TeA** | 97 | 97 | 1 | 8 | 8.2 | 70 | 1 | 6 | 8.6 |
| | | | 2/3 | 27 | 27.8 | | 2/3 | 26 | 37.1 |
| | | | 4 | 17 | 17.5 | | 4 | 14 | 20 |
| | | | 5 | 45 | 46.4 | | 5 | 24 | 34.3 |
| | | | 6 | - | - | | 6 | - | - |
| | 96 | 96 | 1 | 13 | 13.5 | 80 | 1 | 12 | 15 |
| | | | 2/3 | 41 | 42.7 | | 2/3 | 38 | 47.5 |
| | | | 4 | 12 | 12.5 | | 4 | 14 | 17.5 |
| | | | 5 | 30 | 31.3 | | 5 | 16 | 20 |
| | | | 6 | - | - | | 6 | - | - |

### Table S3 – MU pure tone response statistics

| | Spont. FR [Hz] | Bandwidth [#num] | Lifetime sparseness | BF [kHz] | Evoked FR [Hz] | Minimal latency [msec] | Latency to peak [msec] |
|---|---|---|---|---|---|---|---|
| **AUDp** | 1.4 (0.3-3.2) | 9 (1-21) | 0.61 (0.36-0.80) | 16.4 (7.9-28.9) | 19.6 (12.2-32.8) | 34 (23-61) | 98 (22-148) |
| **AUDv** | 0.7 (0.2-2.3) | 8 (0-20) | 0.68 (0.49-0.85) | 16.4 (10.4-35.6) | 19.6 (11.4-33.1) | 40 (27-80) | 100 (25-142) |
| **TeA** | 0.3 (0.2-1.2) | 6 (0-21) | 0.72 (0.52-0.80) | 13.1 (7.6-63.8) | 19.6 (13.1-40.0) | 42 (28-60) | 114 (32-154) |

### Table S4 – Anesthetized data response statistics

| | **Pure Tone protocol** | | | | |
|---|---|---|---|---|---|
| | #Num recorded units | Units per layer | | Percent of total units [%] | Responding to PTs |
| **AUDp** | 151 | 1 | - | - | 150 (99%) |
| | | 2/3 | - | - | |
| | | 4 | 3 | 2 | |
| | | 5 | 117 | 77 | |
| | | 6 | 31 | 21 | |
| **TeA** | 48 | 1 | - | - | 47 (98%) |
| | | 2/3 | 6 | 12.5 | |
| | | 4 | 16 | 33 | |
| | | 5 | 21 | 44 | |
| | | 6 | 5 | 10.5 | |
